# MYC amplifies mitotic perturbations elicited by LXY18 to enable synthetic lethality

**DOI:** 10.1101/2023.11.07.565938

**Authors:** Julia Kalashova, Chenglu Yang, Yan Long, Hongmei Li, Shenqiu Zhang, Ting Zhang, Duo Yu, Xumei Liu, Gang Lv, Namrta Choudhry, Hong Liu, Qiong Shi, Dun Yang

## Abstract

The MYC oncoprotein represents an intriguing target for cancer treatment, but its therapeutic potential has been hindered by the absence of specific pharmacological inhibitors. In this study, we demonstrate that the phenoxy quinoline compound LXY18 selectively targets and eliminates cells overexpressing MYC, leaving non-transformed cells unharmed. This synthetic lethality arises from an acute induction of multipolarity, resulting in a persistent arrest in early mitosis followed by cell death in mitosis or after mitotic slippage. Distinctively, LXY18’s action contrasts with other antimitotic compounds, as they either fail to induce mitotic arrest or elicit mitotic arrest irrespective of MYC abundance. Furthermore, the MYC abundance in a panel of 98 tumor cell lines correlates with their sensitivity to LXY18. Collectively, our findings uncover LXY18 as an MYC- enabled mitotic blocker and open a new avenue to selectively target MYC-overexpressing tumor cells without affecting normal cells.

## Introduction

The *MYC* family of proto-oncogenes represents the most frequently deregulated oncogenic components in human malignancies^1, 2^. Initially cloned as a homolog of v-MYC, the avian viral myelocytomatosis oncogene^3^, MYC serves as the founding member of this family, which also encompasses MYCN (for neuroblastoma-derived homolog) and L-MYC (lung carcinoma-derived homolog)^4^. In this study, all three members are collectively referred to as MYC unless otherwise stated. MYC encodes a basic region/helix-loop-helix/leucine zipper (bHLHZip) protein (MYC) that acts as both a sequence-specific and general transcription factor, regulating > 10% of the human genome, ^5^. To function in this capacity, MYC heterodimerizes exclusively with MAX, another bHLHZip transcription factor^2^. Overexpressed MYC not only promotes uncontrolled proliferation of tumor cells but also offers immune invasion by upregulating immune checkpoint proteins PDL1 and the other do-not-eat me signals^4^.

The goal of disabling MYC has remained central to oncology drug development^6^. However, this objective has proven challenging due to the difficulty in directly inhibiting MYC with small-molecule compounds. This challenge stems from the inherent complexity of targeting transcription factors. Moreover, MYC’s involvement in a multitude of vital physiological functions leads to apprehension regarding the risk of severe side effects following inhibition. A clinically viable therapy that inhibits MYC directly would, therefore, might necessitate meticulous tuning of the treatment dose and duration. Complicating matters further, many cancer cells with abundant MYC do not perish upon MYC depletion, pointing to compensatory pathways that could sustain cancer cell survival. To circumvent these challenges, researchers are exploring alternative therapies that create synthetic lethal (SL) interactions with MYC overexpression, achieving selectively killing of cancer cells while sparing normal cells^7^.

Through its transcriptional activity, overexpressed MYC induces dependencies on genes governing diverse cellular processes such as transcription, splicing, metabolism, mitosis, and apoptosis^7^. During mitosis, certain MYC target genes safeguard the integrity of centrosomes, the mitotic spindle, and kinetochores^8–10^. Consequently, an array of mitotic abnormalities arises upon MYC overexpression, such as excessive asters, altered spindle morphology, and delayed satisfaction of the spindle assembly checkpoint. Interestingly, these abnormalities do not block cells in mitosis and even does not extend the overall time that a cancer cell spend in mitosis^9, 11^. Nevertheless, overexpressed MYC magnifies mitotic disruptions triggered by various antimitotic agents, yielding synthetic lethality. Identified MYC-SL mitotic targets include Polo-like kinase 1 (PLK1), Aurora kinase A and B (AURKA and AURKB), TPX2, KIF11/EG5, each of which is also positively regulated by MYC or could stabilize MYC^7–9, 11, 12^.

In paradoxical to its role in promoting tumorigenesis, overexpressed MYC enhances apoptosis in response to diverse stresses by altering the balance between pro-apoptotic and anti-apoptotic factors to favor cell death^8, 13^. This shift in balance reveals an innate vulnerability that could be exploited by therapies that amplify MYC-primed mitotic irregularities. The multifaceted interplay between MYC overexpression, apoptosis sensitization, and perturbations in mitotic processes highlights the complexity of MYC’s functions and its potential as a SL target for effective therapeutic strategies. These insights underscore the prospect of employing MYC’s own regulatory mechanisms against it, crafting a pathway towards more effective and selective interventions in the ongoing battle against MYC-overexpressing cancers.

Identified through screening for AURKB relocation blockers, (N-(3-((6- bromoquinolin-4-yl)oxy)-5-methoxyphenyl) acetamide (LXY18) has emerged as a promising development candidate^14^, displaying the pharmacokinetic properties suitable for oral administration, sufficient blood-brain barrier penetration, and selective lethality towards cancer cells but spare dividing normal cells^15, 16^. LXY18’s actions extend beyond merely obstructing the mitotic relocation of AURKB in late mitosis. LXY18 also provokes sustained prometaphase arrest selectively in cancer cells but not nontransformed cells and elicits a growth-suppressive effect pattern across diverse cancer cell lines distinct from that associated with AURKB kinase inhibitors^16^. Moreover, some cancer cell lines refractory to AURKB inhibitors are sensitive to LXY18^14^. Despite this progress, the mechanism underpinning the cancer-selective mitotic blocking activity of LXY18 is still a mystery. Nevertheless, the divergence in action, coupled with its ability to induce mitotic catastrophe selectively in cancer cells, makes LXY18 a novel therapeutic warranting further investigation.

In this study, we reveal a positive correlation between MYC abundance and LXY18 sensitivity across 98 human cancer cell lines. Ectopic expression of MYC confer cells sensitivity to LXY18. MYC primes cells for mitotic vulnerability, enabling a SL interaction with LXY18. Specifically, when MYC is overexpressed, LXY18 triggers a prolonged arrest in prometaphase, preceded by the rapid induction of supernumerary centrosomes and excessive asters, which are AURKA-dependent and independent, respectively. This mitotic arrest culminates in apoptosis, eliminating most cells. A minority escapes death in mitosis but only to perish postmitotic slippage or become multinucleated. These insights contribute to our understanding of how LXY18 selectively targets cancer cells as opposed to dividing normal cells. Our findings reveal the potential of LXY18 as a novel MYC-enhanced antimitotic agent for treating MYC-overexpressing cancers, which lack targeted therapies.

## Material and Methods

### Cell culture and chemicals

Chemicals, cell lines, and their culture procedure were described in supplemental information. Each line was amplified and stored in aliquots in a liquid nitrogen tank when it was commercially acquired. Each aliquot was used only for 10-15 passages to minimize the risk of genetic drift, ensuring consistency in the experimental data. Mycoplasma contamination of each cell line was regularly evaluated during the duration of the project by using a PCR-based Myco-LumiTM Luminescent Mycoplasma Detection Kit (Beyotime, Cat. No. C0298M).

### Time-lapse video recording analysis of cell division and cell death

The live-cell imaging experiments were conducted as described before^16^.

### CTG, LDH, and Trypan blue exclusion assays

The Cell-titer Gro (CTG) assay for the live cell number and lactate dehydrogenase (LDH) assay for dead cells were utilized to generate drug concentration-response curves as described in supplemental information. The normalized data were then fit to the four parameters logistic curve using GraphPad Prism 8.0.2. Statistical analysis was performed using the same software. Data are presented as mean ± SD. The Trypan blue exclusion assay uses Trypan blue dye solution at a final concentration of 0.2% to selectively stain dead cells as described^16^. The fraction of dead cells was calculated by dividing the number of dead cells by the total number of cells, both fractions were enumerated under an inverted tissue culture microscopy.

### Cell synchronization

Cells were synchronized at the G1/S boundary by exposure to 2 .5 mM of thymidine for 24 h before released for 16 h into a thymidine-free medium containing 5 μM of CDK1 inhibitor RO3306 that arrested cells at the G2/M transition. The cells were then released into a drug-free medium containing different mitotic inhibitors for 30 min for experiments in Figure 4 and Figure 6. Or cells were released into a fresh drug-free medium for 55 min, then exposed to the drugs 10 min or 30min for experiments in Figure 7 before being fixed for immunofluorescent analysis.

### Immunofluorescent analysis and Western blot

Analysis of protein expression by Western blot and protein localization with immunofluorescent analysis were described^16^ and detailed in the supplemental information.

### Statistical analysis

The dichotomy thresholds for MYC and AUC were chosen to optimize the aggregate values of sensitivity and specificity. This optimization was achieved using the cutpointr tool (version 1.1.2) within the R software environment (version 4.1.2).

Fisher’s exact test p values are calculated with the fisher.test function of the stats package in R version 4.1.2. Cell line mutation data were downloaded from depmap.org. Data on MYC family gene amplification were sourced from the CCLE Broad 2019 dataset at cBioPortal (https://www.cbioportal.org/).

## Results

### A positive correlation between MYC and LXY18 sensitivity in human cancer cell lines

To identify oncogenic driver mutations that might confer sensitivity to LXY18, growth inhibition experiments were conducted in a panel of 98 human cancer cell lines of various origins (Figure 1A and Table S1). ATP levels was assayed as a surrogate for the live cell number to generate drug concentration-response curves (Figure 1B). LXY18 showed potent cytotoxicity, with half-maximum inhibitory concentration (IC_50_) less than 1000, 100, and 10 nM in 81(83%), 76 (78%), and 4 (4%) out of the 98 cancer cell lines, respectively (Figure 1C). Despite of the wide-spectrum activity, the responses to LXY18 varied dramatically from most sensitive in NCI-H446 (IC_50_ = 7 nM) to intrinsically insensitive in NCI-H2170 (IC_50_ >1 μM) (Figure 1C). This heterogeneity indicates that genetic or epigenetic alterations in these cell lines might determine their sensitivity to LXY18.

**Figure 1.**
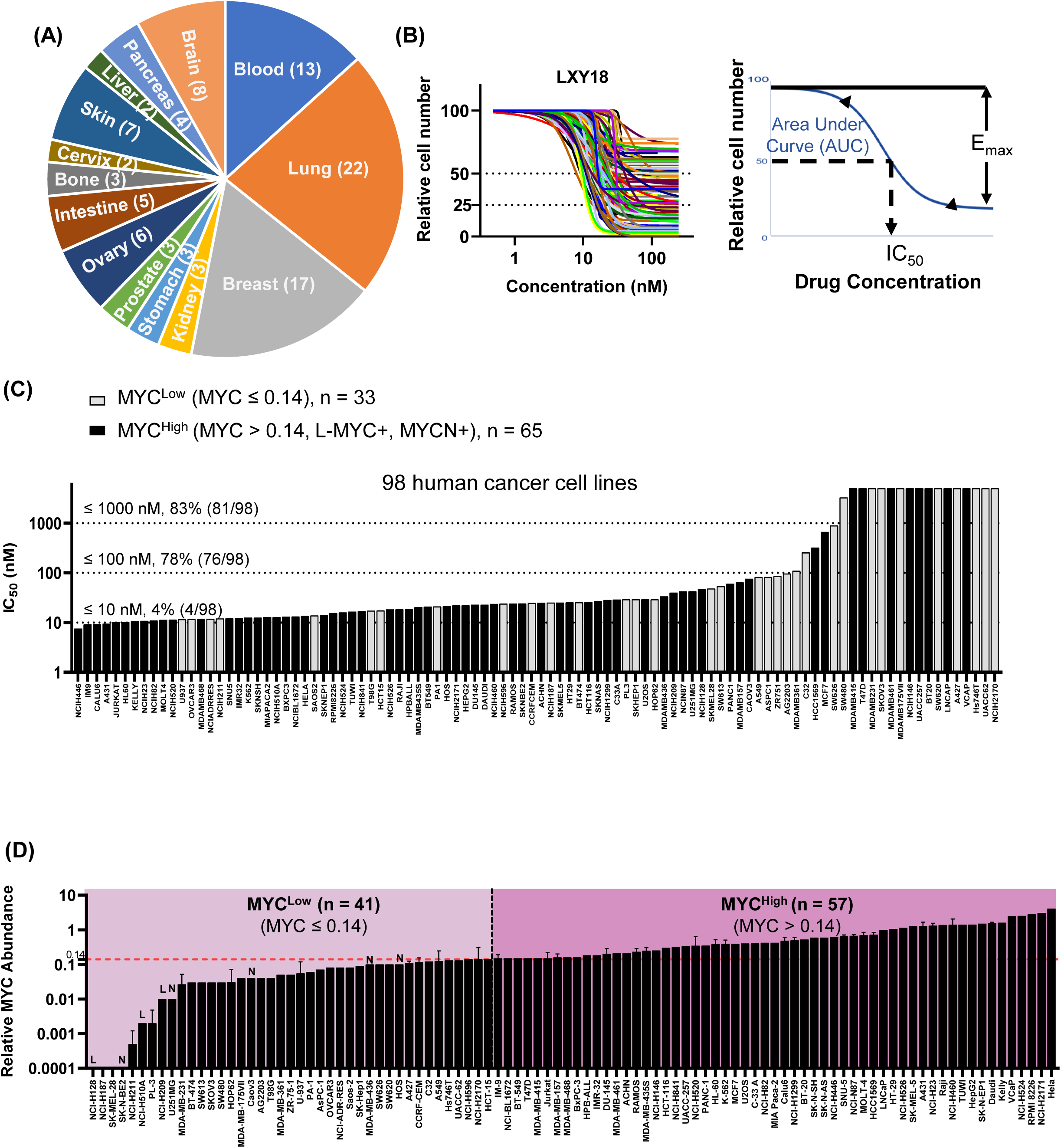
The IC_50_ for LXY18 and MYC abundance in a panel of 98 cancer cell lines. (A) The tissue origins for 98 human cancer cell lines. (B) The response curve of LXY18 in 98 cancer cell lines. Each cell line was treated with LXY18 for 3 days at ten concentrations and then the cell number was determined by the CTG assay and normalized to that of cells treated with vehicle to generate drug concentration-response curves. (C) The IC_50_ of LXY18 in 98 human cancer cell lines. IC_50_ was then calculated by GraphPad Prism 8.0.2. (D) The expression of the MYC family oncoprotein in 98 cancer cell lines. The indicated cell lines were analyzed for MYC, L-MYC and MYCN by Western blot. The MYC level in a cancer cell line was normalized to that in RPE-MYC cells, which was set as 1.0. The Y- axis represents the relative MYC abundance. L on the graph marks a cell line positive for L-MYC. N denotes a cell line positive for MYCN. Only cell lines with MYC less than one tenth of that in RPE-MYC cells were analyzed for L-MYC and MYCN. A MYC cutoff of 0.14 was identified to optimally define MYC^Low^ and MYC^High^ cell lines regarding to their response to LXY18.

The status of the tumor suppressor genes *TP*53, *RB*1, and *PTEN*, as well as the presence of amplification of the *MYC* (*MYC*^AMP^) family of oncogenes (*MYC*, *MYCN* and L-*MYC*) and activating mutations for other oncogenes *RAS*, *BRAF*, and *PI3K*, were available for 78-91 of our 98 cell lines (Table S2). *MYC*^AMP^ failed to predict a favorable response to LXY18 (Figure S1A, B). Likewise, none of the other oncogenic driver mutations showed a statistically significant correlation with LXY18 sensitivity when examined either individually or collectively as one pathway (Figure S1C-F).

*MYC*^AMP^ does not necessarily lead to abundant expression of its protein product. Conversely, due to up-regulation of its transcription activators or stabilization at the level of mRNA or protein, abundant MYC frequently occurs in human malignancies that lack *MYC*^AMP 17^. We, therefore, quantified MYC, MYCN, and L-MYC in the panel of cell lines by using the Li-COR NIR fluorescence detection system after Western blot (Figure S2). A continuity in the MYC abundance was observed from undetectable to readily detected and highly abundant (Figure 1D). There were 66 cell lines with abundant MYC not less than one tenth of that in RPE-MYC cells, which have been engineered to express MYC and are highly responsive to multiple MYC-SL compounds^12, 18^. The remaining 32 lines harbored less MYC and were further examined for L-MYC and MYCN. Among them, the small cell lung carcinoma (SCLC) cell line NCI-H510A is known to have amplification and expression of L-MYC^19^. Two more cell lines, NCI-H128 and NCI-H209, were also identified as positive for L-MYC, because they had L-MYC no less than 50% of that in NCI-H510A (Figures 1D and S2B). The neuroblastoma cell line SK-N-BE2 is known to have amplification and overexpression of MYCN^20^. Five lines (MDA-MB-436, CaoV3, HOS, SK-N-BE2, and U251MG) were tested positive for MYCN because four of them had MYCN comparable to that in SK-N-BE2 (Figure 1D and S2C).

The shape of drug concentration-response curves among different cell lines for a given drug could vary significantly, making IC_50_ values not always the most accurate metric for comparing drug potency among different cell lines. We chose to analyze the correlation between the MYC abundance and LXY18 response by using multi-descriptors including IC_50_, the maximum effect (E_max_), and the area under the curve (AUC) (Figure 1B). In the pane of cell lines treated with LXY18, MYC abundance was positively correlated with E_max_ and negatively correlated with IC_50_ and AUC when the threshold of relative MYC abundance was set at 0.14 (Figure 2A-C). AUC was used for further analysis as partitioning its values at 0.73 maximized sensitivity and specificity. This placed 70% (47/67) of responsive cell lines in the MYC^High^ group (Figure 2D). Based on this stratification, 82% of cell lines in the MYC^High^ group and 54% of cell lines in the MYC^Low^ group were sensitive to LXY18 (Figure 2D). The positive correlation between MYC and LXY18 sensitivity held true when multiple cutoffs for MYC expression and AUC were examined (Figure S3).

**Figure 2.**
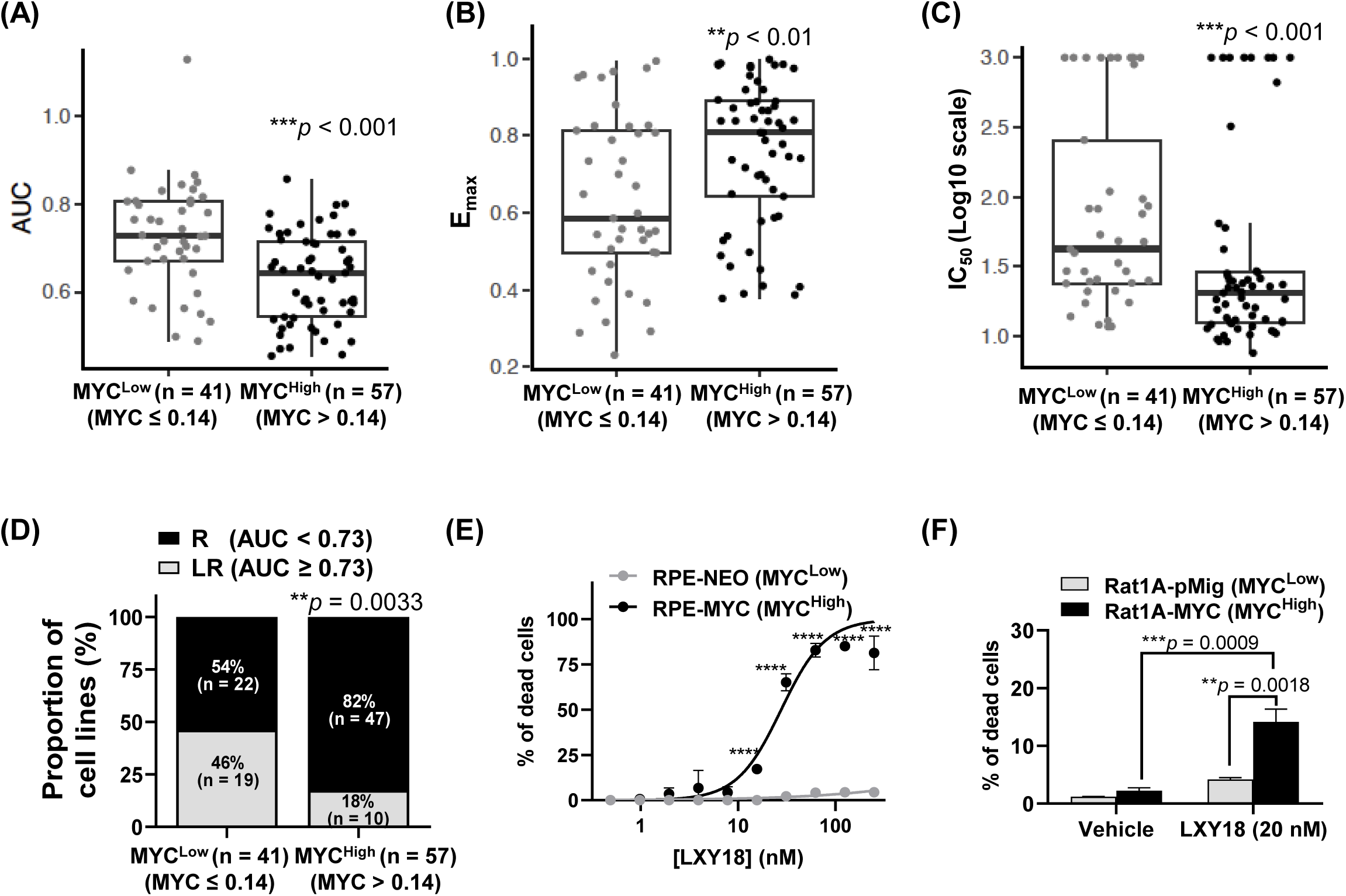
The abundance of MYC dictates sensitivity to LXY18. Correlation analysis of MYC abundance and LXY18 sensitivity was performed with response-drug concentration descriptors AUC (A), Emax (B), and IC_50_ (C) at a MYC cut- off level of 0.14. For each panel, a box plot presentation of cell lines in the MYC^High^ group or MYC^Low^ group. Each dot is a cell line. *p*-values indicate the statistical significance between the two groups. D. The proportion of cell lines that were responsive versus less responsive, as assessed through AUC measurement, that fit into the MYC^High^ and MYC^Low^ groups. Cell lines were stratified based on optimal thresholds of MYC expression and LXY18 sensitivity indicated by AUC. The threshold for MYC and AUC is 0.14 and 0.73, respectively. R: Responsive. LR: Less Responsive. (E) MYC-primed killing of epithelial cells. RPE-NEO and RPE-MYC cell lines were treated with LXY18 or vehicle (0.1% DMSO) for 72 hours at indicated concentrations and then the cell death was determined by the LDH assay. Data, here and ensuing panel, are presented as mean ± SD from three independent experiments. ****p<0.0001. (F) Sensitization of fibroblasts by MYC. Rat1A- pMig and Rat1A-MYC cell lines were treated with 20 nM of LXY18 for 48 hours and then the dead cells were determined by the Trypan blue assay and the percentage of dead cells in the population was calculated. ***p<0.001, **p<0.01.

Despite of this strong correlation between MYC abundance and LXY18 sensitivity, there were exceptions in both groups. Ten out of 57 cell lines in the MYC^High^ group were inert to LXY18 whereas 22 out of 41 cell lines in the MYC^Low^ group were sensitive to LXY18 (Figure 2D). Nevertheless, collectively, these data suggest that the abundance of MYC in the human cancer cell lines positively relates to their sensitivity to LXY18. We reached the same conclusion when all MYC family members were included (Figure S4A- C). Inclusion of all MYC family members in the correlation analysis improved accuracy as indicated by reducing the *p*-value from 0.0033 (Figure 2D) to 0.0003 (Figure S4D).

### Ectopic expression of MYC confers sensitivity to LXY18

To address whether MYC was causative to LXY18 sensitivity, we examine LXY18 responses in two pairs of isogenic cell lines. One pair includes human retinal pigment epithelial cells (RPE), which have been engineered to express a Neomycin-selection marker (RPE-NEO) or the MYC protooncogene (RPE-MYC), respectively^12^. To quantify the cytotoxicity of LXY18, we assayed lactate dehydrogenase (LDH) released into the medium from dead cells. This assay revealed that LXY18 killed more than 65% of RPE-MYC cells but almost none in RPE-NEO cells when used at a concentration ranging from 31 to 250 nM (Figure 2E). Likewise, this MYC-enabled lethality was detected in the other pair of cell lines, which are rat embryo fibroblasts modified to express either MYC (Rat1A- MYC) or an empty pMig vector (Rat1A-pMig)^12^ (Figure 2F). Thus, MYC-LXY18 SL interaction is conserved between epithelial cells and fibroblasts.

### MYC enables the sustained mitotic arrest and lethality elicited by LXY18

To understand the mechanism underpinning MYC-LXY18 SL interaction, a time-lapse video filming experiment was conducted to document the effect of MYC and LXY18 on RPE cells individually or collectively (Figure 3A). Without drug treatment, the mitotic duration increased from 0.85 ± 0.31 to 4.9 ± 3.0 hours when MYC was overexpressed (Figure 3B). Treatment with LXY18 dramatically increased the average duration of mitosis to 11.4 ± 4.9 and 22.1 ± 10.1 hours for RPE-NEO and RPE-MYC cells, respectively (Figure 3B). This synergism between MYC and LXY18 was also apparent when the mitotic index (MI) was evaluated. About 63-84% of RPE-MYC cells remained in mitosis 24 or 48 hours after treatment whereas the MI of RPE-NEO cells were never above 13% in the presence of LXY18 (Figure 3C). This high MI reflected a prolonged mitotic arrest because no cells entered mitosis twice during this experiment (Figure 3A). This filming study also revealed that the mitotic arrest of RPE-NEO cells by LXY18 was transient. All RPE-NEO cells eventually exited from mitosis without completion of cytokinesis, became multinucleated, remained alive, and failed to enter the next mitosis (Figure 3A, D).

**Figure 3.**
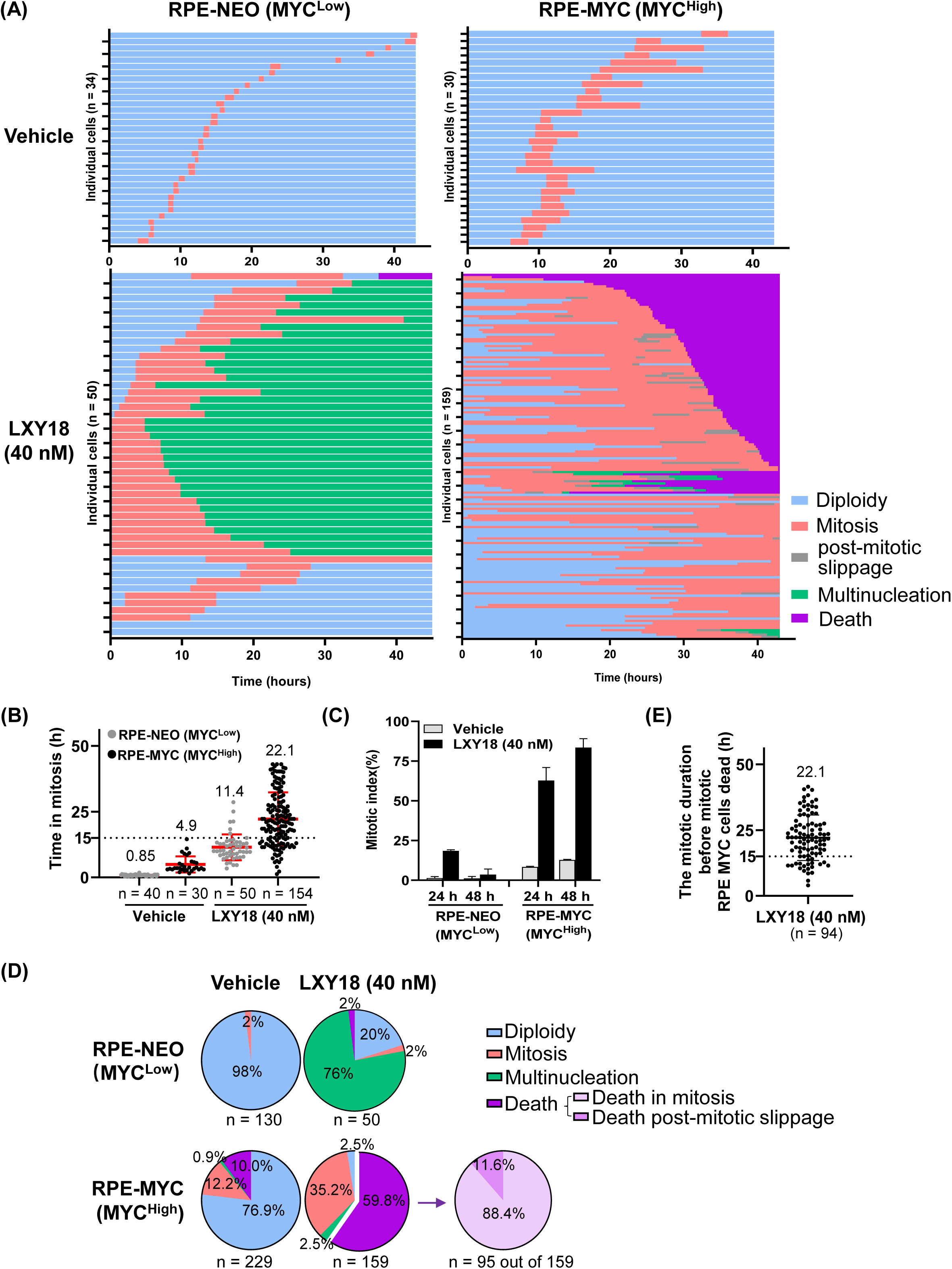
MYC and LXY18 act synergistically to elicit mitotic catastrophe. After being exposed to 0.1% DMSO or 40 nM of LXY18, indicated cells were subjected to time-lapse filming for 48 h. (A) Graphic summary of the fate of individual RPE-NEO and RPE-MYC cells. (B) Synergistic induction of the mitotic arrest by MYC and LXY18. n indicates the number of mitotic cells scored in each group. (C) MYC-primed mitotic blockage. The histogram shows the mitotic index 24 and 48 h after initiating the treatment. 60∼240 cells were scored at each time point in each group. (D) The faction of cells with the indicated status at the endpoint of this filming experiment. n indicates the number of mitotic cells scored in each group. (E) The time an RPE-MYC cell spent in mitosis before dying. Each dot represents a cell.

Consistent with our early findings (Figure 2E), cell death occurred extensively in RPE-MYC but not RPE-NEO cells in the presence of LXY18 (Figure 3A, D). The death was mainly observed during mitosis or upon multipolar division, with less than 12% of death cases appeared postmitotic slippage (Figures 3A, D and S5A). This MYC-primed mitotic death, however, could not be simply explained by longer mitotic duration when MYC was overexpressed. LXY18-treated RPE-NEO cells spent 11.4 ± 4.9 hours in mitosis without induction of cell death whereas 21.3% (20 out of 94) of mitotic death cases occurred after RPE-MYC cells spent less than 15 hours in mitosis (Figure 3B, E). This contrast implied that the proapoptotic activity of MYC was also required for LXY18 to kill cells, consistent with the reports that MYC sensitizes mitotic cells to apoptosis^8^. A small fraction of RPE-MYC cells escaped from the mitotic death by undergoing mitotic slippage or returning to interphase without completion of cytokinesis (Figure 3A). Many of them subsequently became extinct, limiting the accumulation of the polyploid cells (Figures 3A, D and S5A).

In response to LXY18, RPE-MYC cells behaved indistinguishable from cancer cell lines that intrinsically overexpress MYC^16^, undergoing a prolonged mitotic arrest. This MYC-enabled mitotic blockage is not a general phenomenon to any disturbance of mitosis because a variety of other antimitotic agents failed to cooperate with MYC to elevate the MI. In this comparative study, LXY18 provoked robust mitotic arrest only when MYC was overexpressed (Figure 4A, B), achieving an MI ratio of 18.7 (Figure 4C). In contrast, compounds known to drive cells through mitosis even though mitotic abnormalities activate the spindle assembly checkpoint^11^, including an AURKB-specific inhibitor AZD1152, a pan-AURK inhibitor AMG900, and a monopolar spindle 1 (MPS1) inhibitor BAY1217389, all failed to significantly elevate the MI in RPE cells irrespective of MYC abundance (Figure 4B). Likewise, a Polo-like kinase 4 (PLK4) inhibitor (Centrinone) did not change the MI in either of the two cell lines (Figure 4B-C). Furthermore, a centrosome-associated protein E (CENP-E) inhibitor GSK923295 and a kinesin-5/EG5/KIF11 inhibitor SB743921 slightly elevated the MI in both cell lines, with an MI ratio of 1.1 and 1.9 in RPE-MYC cells relative to control RPE-NEO cells, respectively (Figure 4A-C). KIF18A inhibitor Sovilnesibl heightened the MI ratio to 2.6 (Figure 4C). Thus, the MYC-primed mitotic blocking activity of LXY18 is unique and sets it apart from other antimitotic agents.

**Figure 4.**
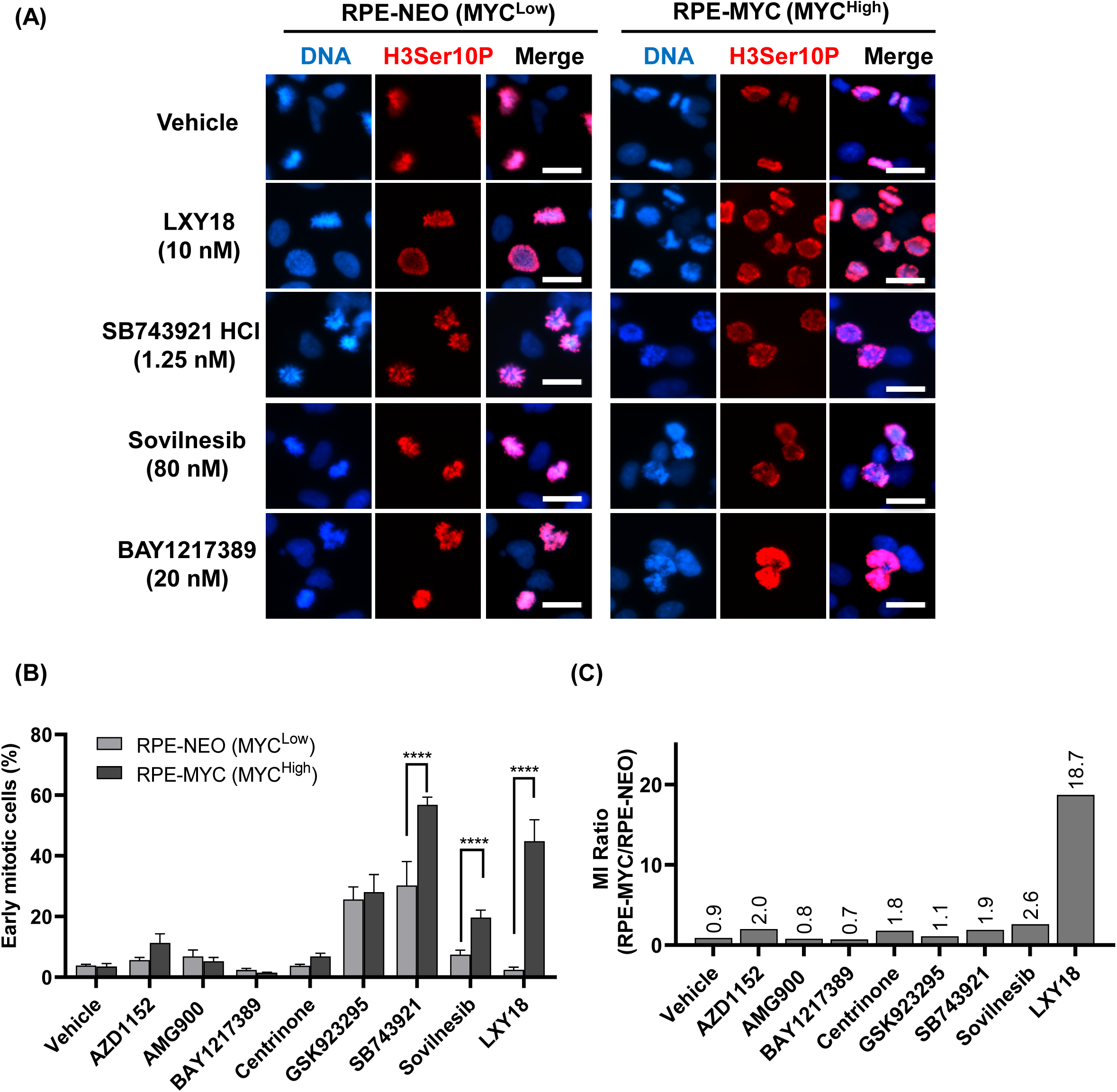
LXY18 but no other antimitotic agents arrest cells in early mitosis when MYC is overexpressed. RPE-NEO and RPE-MYC cells were treated with either vehicle control (0.1% DMSO) or indicated mitotic inhibitors for 24 h before immunofluorescent staining for H3Ser10P and staining for DNA with 4’,6-diamidino-2-phenylindole (DAPI). Cells positive for H3Ser10P were scored as mitotic cells and quantified when cells were treated with compounds other than AURKB inhibitors. For cells exposed to AZD1152 and AMG900, mitotic cells were identified based on their condensed chromosomes revealed by DAPI staining. The following mitotic inhibitors were used at minimally effective concentrations that are known to inhibit their respective targets, AZD1152 (200 nM), AMG900 (10 nM), BAY1217389 (20 nM), Centrinone (1 μM), GSK923295 (40 nM), SB743921 HCl (1.25 nM), Sovilnesib (80 nM). (A) Representative immunofluorescent images. (B) The percentage of the mitotic cells. Data are presented as mean ± SD. ****p<0.0001. (C) Histograms show the ratio of the mitotic index (MI) in RPE-MYC cells relative to RPE- NEO cells.

### LXY18 and MYC collaboratively trigger multipolarity

To dissect the mechanism underpinning the MYC-primed mitotic arrest, we analyzed chromosomes in fixed cells after DAPI staining. Treatment of RPE-MYC for 6 hours with LXY18 resulted in an arrest of mitotic cells in prometaphase, as indicated by a lack of apparent chromosome alignment and an identifiable metaphase plate (Figure 5A, B, DAPI staining). In vehicle-treated cells, metaphase cells were readily visible. In accord with chromosome behavior defect, immunofluorescent staining for β-tubulin revealed abnormalities in the mitotic spindle, which provides the mechanical force to move chromosomes. The dominant abnormality was the appearance of ectopic asters in 50-70% of cells treated with LXY18 (Figure 5C). A minor fraction of cells with monopolar spindles was also noted (Figure 5B, C). In contrast, monopolarity and multipolarity were negligible in cells treated with vehicle (Figure 5C).

**Figure 5.**
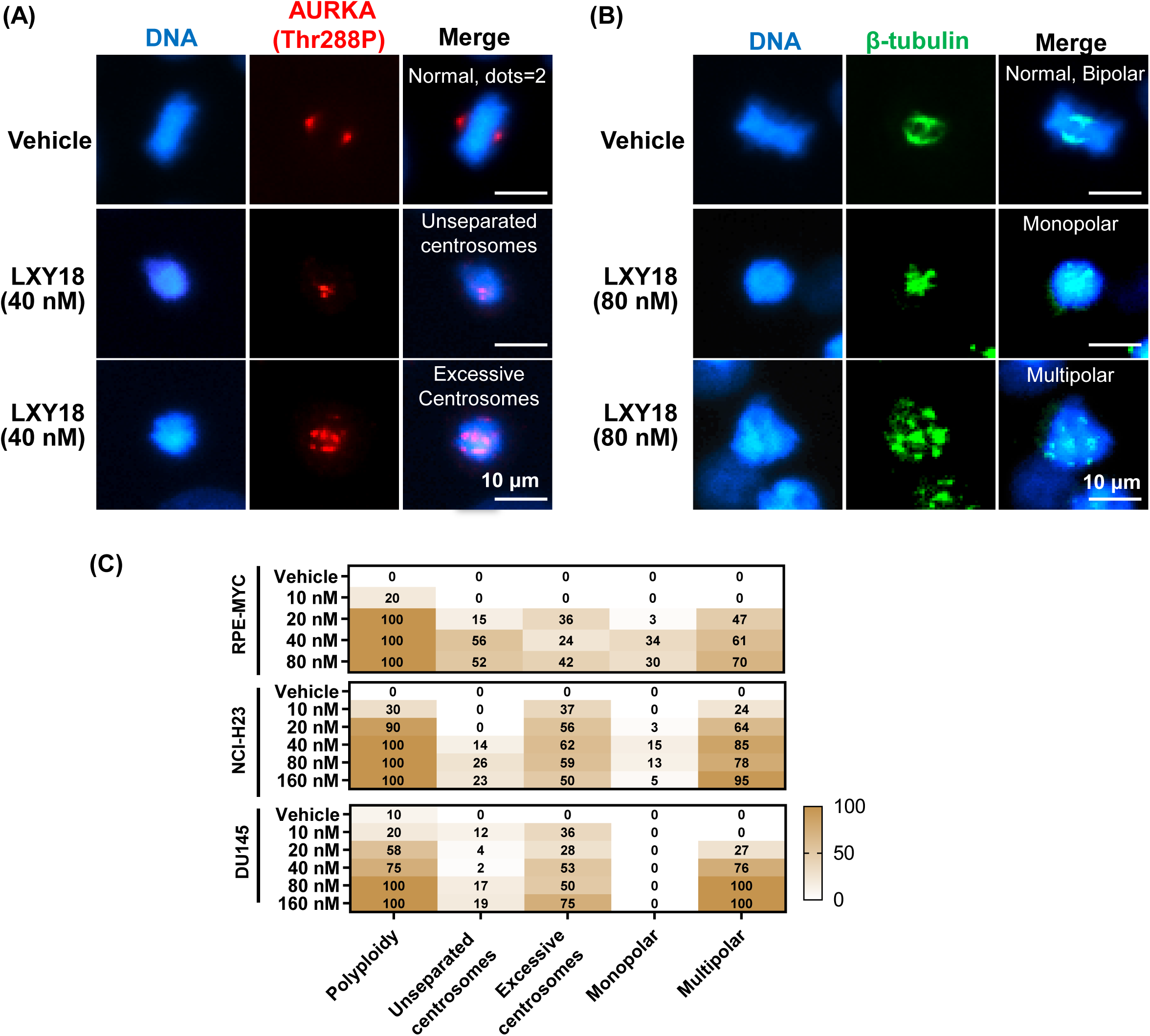
LXY18 acutely elicits ectopic asters and centrosomes in cells with abundant MYC. RPE-MYC, NCI-H23 and DU145 cell lines were treated with either 0.1% DMSO or LXY18 for 6 hours before β-tubulin and AURKA Thr288P were analyzed by the immunofluorescent analysis. Scale bars: 10 μm. (A) Induction of centrosomal abnormalities by LXY18 in RPE-MYC cells. (B) Induction of spindle abnormalities in RPE-MYC cells. (C). The percentage of LXY18 phenotypes in the indicated cell lines.

We reproduced these MYC-enabled spindle defects with cells released from an arrest at the G2/M transition (Figure 6A). Ectopic asters appeared within 30 min after exposing RPE-MYC cells to LXY18 (Figure 6B-C). Under the same treatment condition, LXY18 failed to elicit ectopic asters in RPE-NEO cells. Thus, MYC primes cells to acute induction of ectopic asters by LXY18. Excessive asters, in turn, lead to the sustained activation of the spindle assembly checkpoint response, mitotic arrest and cell death.

**Figure 6.**
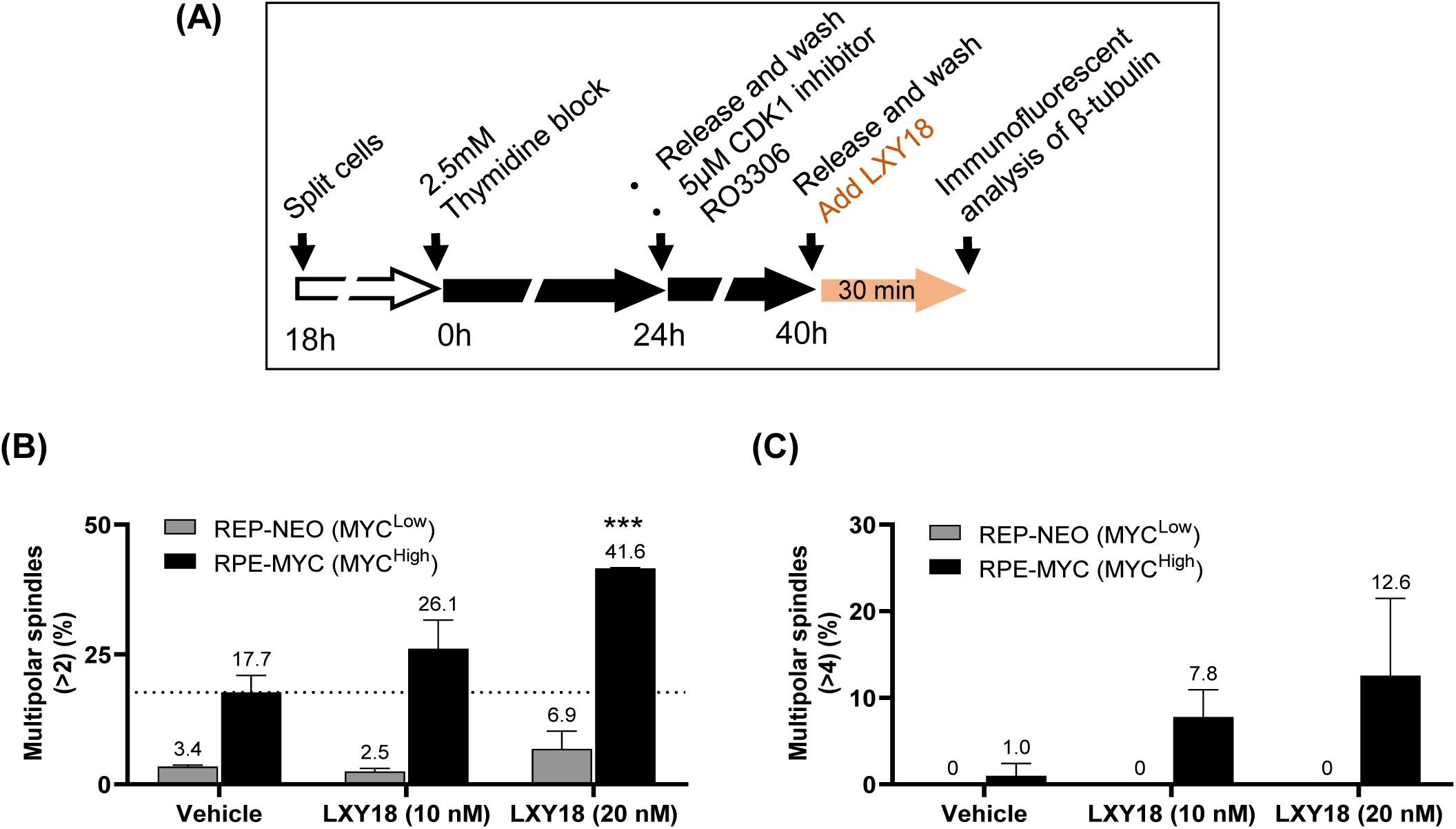
Overexpression of MYC synergizes with LXY18 to acutely elicit multipolarity. The indicated cells were first blocked in S and then the G2/M transition. Upon released for 30 min from the G2/M arrest, cells were treated with either vehicle control (0.1% DMSO) or LXY18 for 30 min before fixed for immunofluorescent analysis for β- tubulin. (A) The flow chart shows synchronization and treatment scheme. (B) Quantification of cells with > 2 asters in the presence of the indicated compounds. (C) Quantification of cells with > 4 asters in the presence of the indicated compounds. Data are presented as mean ± SD. ****p<0.0001, indicate the statistical significance between the LXY18 and vehicle group in each cell line.

### Acute induction of supernumerary centrosomes

Aster microtubules usually nucleate from centrosomes despite that acentrosomal asters could also materialize under a few situations^21, 22^. Induction of ectopic asters by LXY18 prompted us to examine the number and integrity of centrosomes. Immunofluorescent staining was conducted for two components of the pericentriolar material, γ-tubulin and the active AURKA, AURKA Thr288P, which is a marker for its kinase activity and mature mitotic centrosomes^23^. Results with both markers consistently showed that treatment of RPE-MYC cells with LXY18 for 6 hours increased centrosomes in mitotic cells, ranging from three to a dozen (Figure 5A, C). The percentage of mitotic cells with supernumerary centrosomes approximated that of mitotic cells harboring excessive asters (Figure 5C). Both types of abnormalities increased coordinately when the concentration of LXY18 was elevated (Figure 5C). In RPE-MYC cells treated with DMSO, there were always two dots positive for γ-tubulin or β-tubulin. LXY18 also acutely elicited excessive centrosomes and asters in two MYC^High^ human cancer cell lines, a lung cancer cell line, NCI-H23, and a prostate cancer cell line, DU-145 (Figure 5C).

Centrosome replication normally occurs coincidently with DNA replication in the S phase^24^. Double immunofluorescent staining analysis of NCI-H23 cells revealed that all G2 cells, identified by an anti-Cyclin B1 antibody, showed only two γ-tubulin-positive dots when treated for 6 hours with LXY18 (Figure S6). To provide additional evidence that ectopic centrosomes were induced during mitosis rather than in the S phase, we treated NCI-H23 cells released from a prior synchronization at the G2/M transition (Figure 7A). The synchronization process *per se* markedly increased the basal levels of mitotic abnormalities; 30% of mitotic cells displaying ectopic dots positive for either AURKA Thr288P or γ-tubulin (Figure 7B, D) and approximately 20% of mitotic cells also showed ectopic asters (Figure 7C, D). The basal levels for either of the two types of abnormalities in asynchronized mitotic cells was less than 1% (Figure 5). Despite of the high basal abnormalities triggered by the synchronization procedure, the fraction of mitotic cells with ectopic centrosomes and asters further increased to 47 ± 2.8% and 56 ± 2.8% (Figure 7B-D), respectively, after treatment for 10-30 min with LXY18. No excessive centrosomes or asters were found in interphase cells. Collectively, these results suggest that LXY18 acutely induces the formation of ectopic centrosomes and asters.

**Figure 7.**
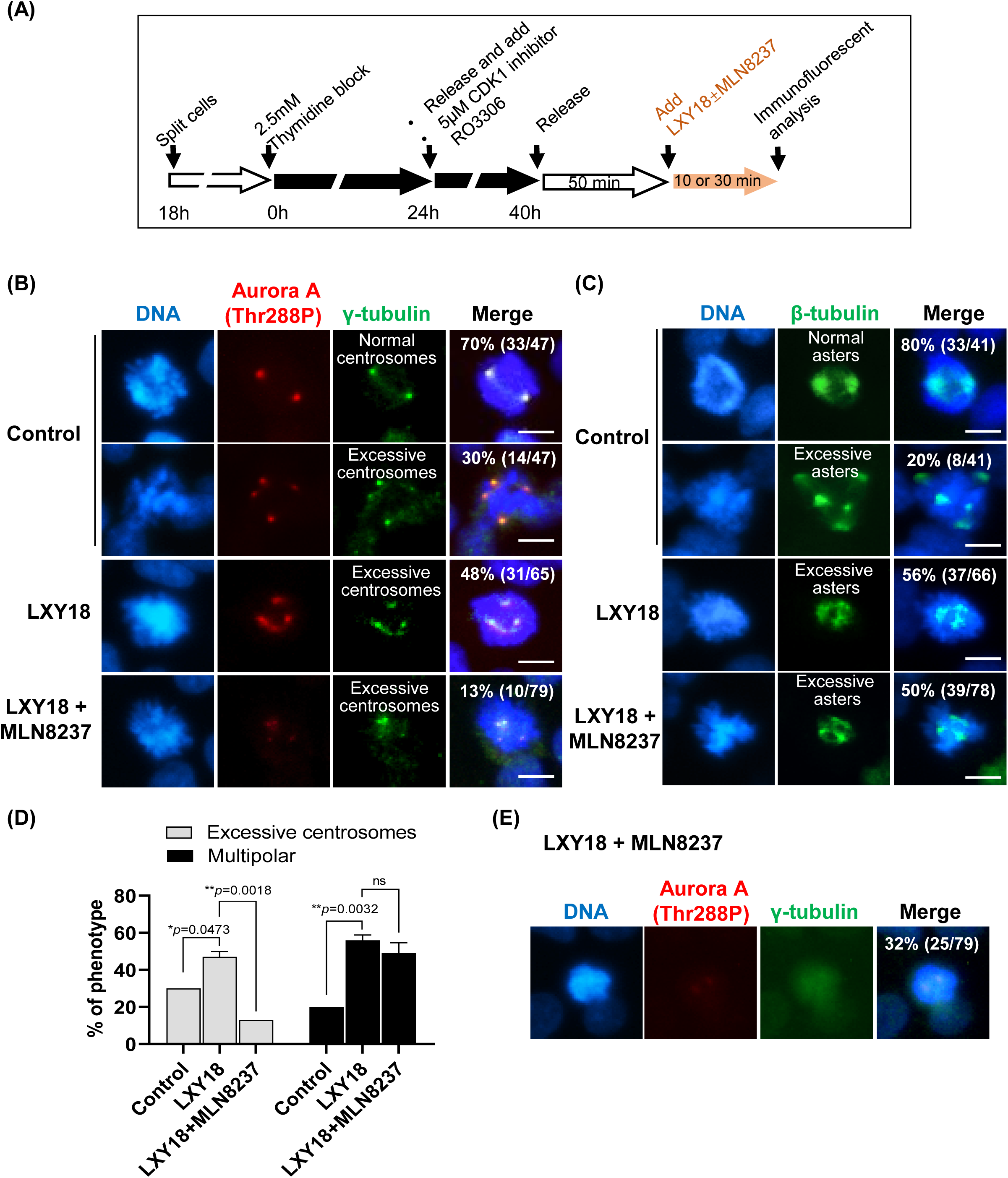
Induction of ectopic centrosomes but not multipolarity by LXY18 requires AURKA. NCI-H23 cells were synchronized, released, and then treated with either vehicle control (0.1% DMSO) or indicated compounds for 10- or 30-min. Bars: 10μm. (A). The flow chart in shows the synchronization procedure. (B, E) Double immunofluorescent staining for γ-tubulin and AURKA Thr288P. (C) Immunofluorescent staining for β-tubulin. (D) The percentage of cells with indicated mitotic abnormalities. Centrosomes and asters were identified by staining for γ-tubulin and β-tubulin, respectively. Combined data at 10 and 30 min were presented because no difference was observed between the two time points. Data are presented as mean ± SD. *p<0.05, **p<0.01.

### AURKA is required to induce excessive centrosomes but not asters

The presence of ectopic centrosomes positive for the active AURKA raised the possibility that induction of ectopic γ-tubulin dots requires AURKA kinase activity. NCI-H23 cells were treated for 10 min with LXY18 and an AURKA-specific inhibitor MLN8237^25^ either individually or in combination. MLN8237 effectively inactivated AURKA, as judged by the diminished signal of the autophosphorylation of AURKA at Thr288 (Figure 7E). The treatment decreased the frequency of cells with ectopic γ-tubulin dots from 48% to 13% (Figure 7B, D). However, the percentage of mitotic cells with ectopic asters only slightly reduced from 56% to 50% (Figure 7C, D). This contrast indicates that induction of ectopic centrosomes but not asters was dependent on the kinase activity of AURKA. LXY18 could induce acentrosomal asters when AURKA was inhibited.

## Discussion

In this study, we used model cell lines and a large panel of human cancer cell lines to demonstrate that ectopically expressed MYC primes cells to the lethal effect of LXY18 and that the MYC abundance is useful to predict LXY18 sensitivity. The MYC-enabled lethality in response to LXY18 could be attributed to their collaborative induction of multipolarity, leading to a prolonged prometaphase arrest and subsequent mitotic catastrophe. Both the mitosis-disturbing activity and the proapoptotic activity of MYC are required to materialize the lethal activity of LXY18. These findings mark the first report of a MYC-primed small-molecule mitotic blocker, showcasing the selectivity of LXY18 in amplifying MYC-induced mitotic vulnerabilities. This study also uncovers a new therapeutic strategy for selectively killing of MYC-overexpressing tumor cells as opposed to dividing normal cells, which have undetectable levels of MYC.

### MYC as a critical determinant of LXY18 sensitivity

MYC is a crucial determinant in the sensitivity of cancer cells to LXY18, despite the extensive genetic variability among different cell lines. This sensitivity is positively correlated with the endogenous MYC in cancer cell lines and is conferred by ectopic MYC in nontransformed cells with otherwise undetectable MYC. Notably, oncogenic alterations other than deregulated *MYC* fail to predict sensitivity to LXY18.

Exceptions, however, exist that defy the MYC-LXY18 SL interaction. These exceptions highlight the need for multiparameter biomarkers centered on MYC to more accurately predict LXY18 sensitivity. The transcriptional activity of MYC is modulated by its antagonists, such as MAD, MXD1 and MIZ, and the availability of its binding partner MAX^26, 27^. Alterations in these factors can influence the transcriptional response to MYC, consequently affecting response to LXY18. In line of this rationale, a MYC target gene signature might provide more accurate reflection of its activity than MYC abundance. However, this approach has intrinsic limitation because MYC regulates more than 10% of the human genome and the specific MYC transcriptional target that confers LXY18 sensitivity is unknown. Nevertheless, the widespread expression of MYC in human malignancies implies a wide-spectrum therapeutic potential for LXY18.

### A biphasic lethality by MYC and LXY18 interaction

The MYC-LXY18 SL interaction is executed by death both in mitosis and post-mitotic slippage. Initially, cells with abundant MYC experience persistent activation of the mitotic spindle assembly checkpoint, leading to mitotic catastrophe. Subsequently, cells that escape this early phase of death undergo MYC-enabled death after mitotic slippage or cytokinetic failure. Yet, a small fraction of cells evades this biphasic death and develops polyploidy. In contrast, MYC^Low^ cells avoid the biphasic death altogether, entering proliferative arrest after incompletion of cytokinesis. The exact contribution of each of the two phases of cell death in MYC^High^ cells depends on the cellular context, with variations likely related to the robustness of the spindle checkpoint response^28^ and expression of the proapoptotic factors such as BCL2 and BCL-xL^29^. This highlights the complexity of MYC-LXY18 SL interaction, the promising virtue of bimodal killing by a single therapeutic agent, and selectivity in killing of cancer cells while sparing normal cells.

### LXY18 as a MYC-primed mitotic blocker

The search for tumor-specific mitotic regulators as oncology targets has long been a goal in cancer research. Despite extensive research, no such a target has been discovered, as identified mitotic regulators control cell division in both normal and cancerous cells without much discrimination. Intriguingly, we have identified LXY18 as a MYC-primed mitotic blocker that acutely triggers multipolarity and a prolonged mitotic arrest when MYC is overexpressed. No other antimitotic agents examined in this study has such a MYC-enabled activity.

LXY18 acutely amplifies the mitotic disturbances in centrosome and spindle integrity elicited by MYC. The LXY18 target, disablement of which elicits MYC-primed mitotic abnormalities, is still unknown. LXY18 does not inhibit kinase activity of the known MYC-SL targets, such as CDK1, PLK1, KIF11/EG5, AURKA, AURKB, and PIM1^16^. LXY18 mislocalizes AURKB in late mitosis, presumably disabling its cytokinetic function. But the sustained mitotic arrest in cells exposed to LXY18 but not AURKB kinase inhibitors indicates that LXY18 must also disable a mitotic regulator other than AURKB.

LXY18 treatment recapitulates multiple defects seen after inhibition of MKLP2, including AURKB relocation failure, the mitotic arrest, and multipolarity^16, 30, 31^. These similarities point to the possibility of MKLP2 as a target for LXY18. However, it is not clear how LXY18 could inhibit MKLP2 since the compound does not inhibit the ATPase activity of MKLP2 or suppress its expression^16^. Regardless, ectopic centrosomes and asters may serve as valuable pharmacological biomarkers, aiding in precise drug dosing of LXY18 for patients and validation of its mechanism of action *in vivo*.

## Acknowledgments

We are grateful to all members of the discovery oncology division of Anticancer Bioscience for their critical review and constructive suggestions.

## Availability of data and materials statement

All data analyzed during this study are included within the manuscript (and its Supplementary Information files). Data deposition does not apply to the current study.

## Supporting information

The supporting information file comprises Figs. S1-6, Tables S1 and S2 in a single PDF file format.

## Supplemental Information

**Figure S1.**
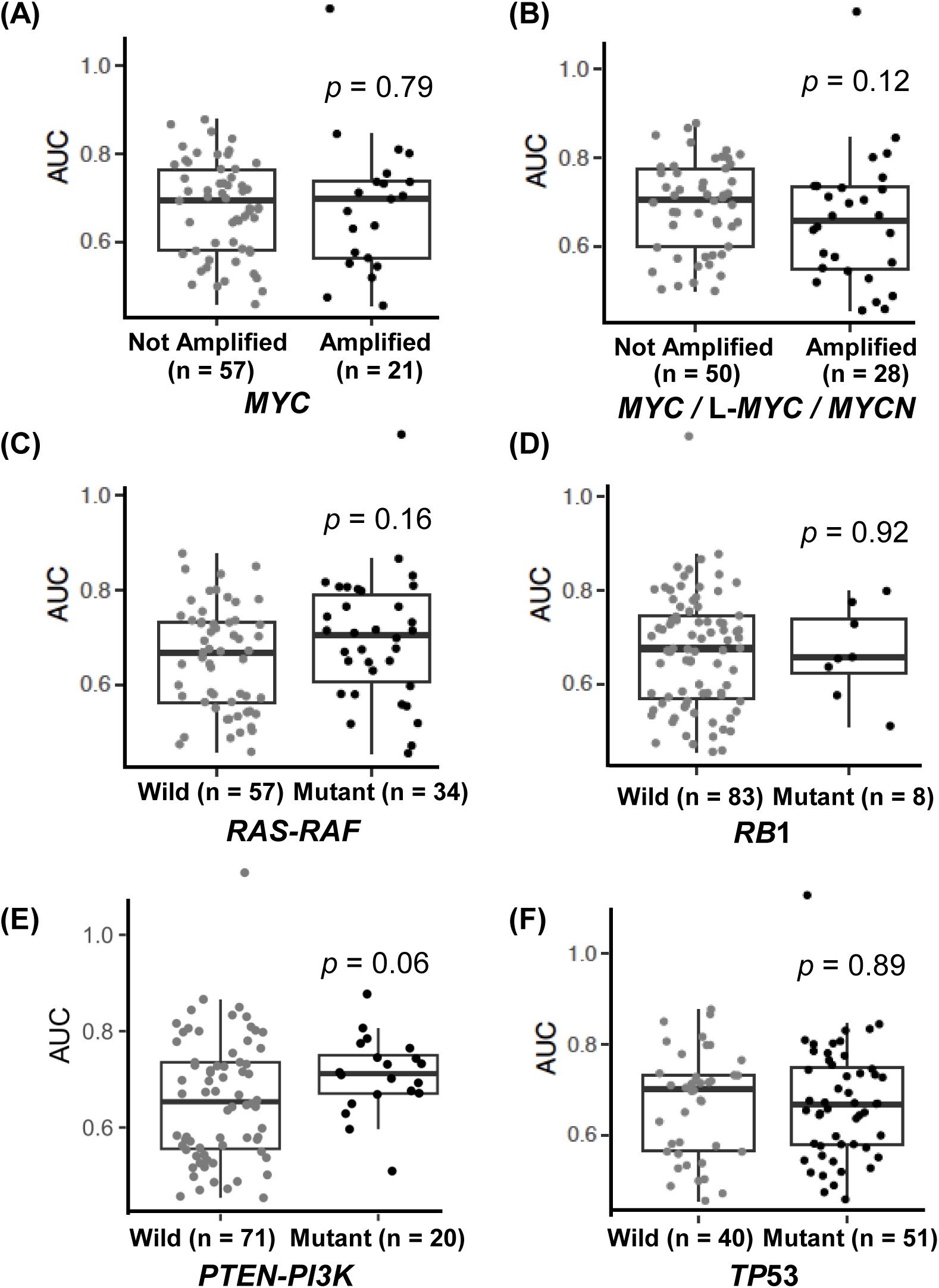
*MYC* amplification and other oncogenic driver mutations fail to predict LXY18 sensitivity. Boxplots show the AUC of each cell line in a define group with specific oncogenic driver mutation. (A) With or without MYC amplification. (B) With or without amplification of any member of the *MYC* family. (C) Wild type or an active mutation in any member of either the *RAS* or *RAF* family. (D) Wild type or mutant *RB*1. (E) Wild type or mutant *PTEN or PI3K*. (F) Wild type or mutant *TP*53.

**Figure S2.**
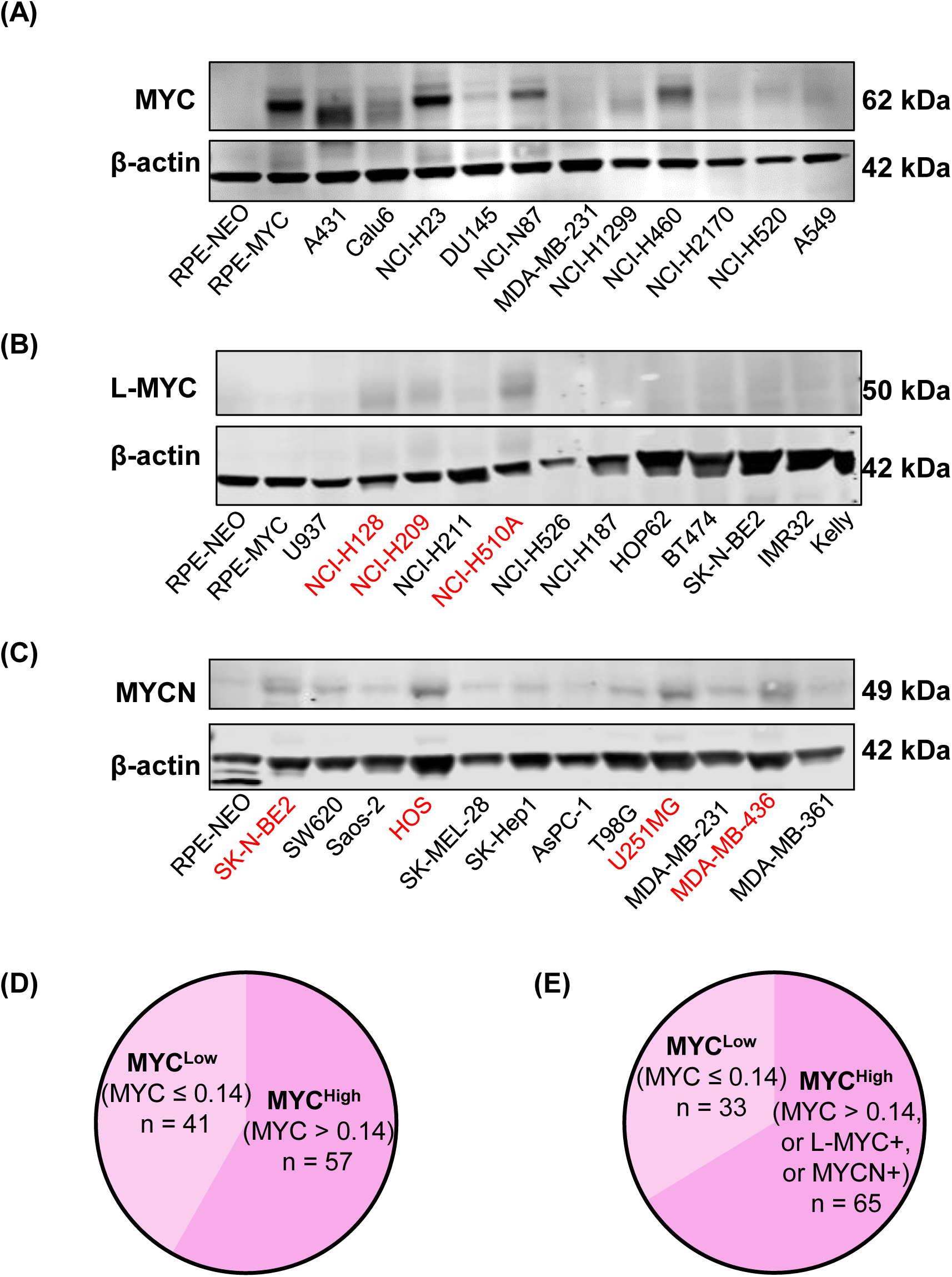
Representative Western blot images of human cancer cell lines. The indicated cell lines were analyzed for MYC (A), L-MYC (B), and MYCN (C) by Western blot. RPE-MYC, NCI-H510A and SK-N-BE2 were used as positive control for MYC, L-MYC, and MYCN, respectively. The data were normalized to these three cell lines. (D) The MYC^Low^ and MYC^High^ groups. The MYC cutoff level is 0.14. (E) The revised MYC^Low^ and MYC^High^ groups. MYC cutoff level is 0.14. MYC^Low^ cell lines positive for L-MYC and MYCN are included into the MYC^High^ group.

**Figure S3.**
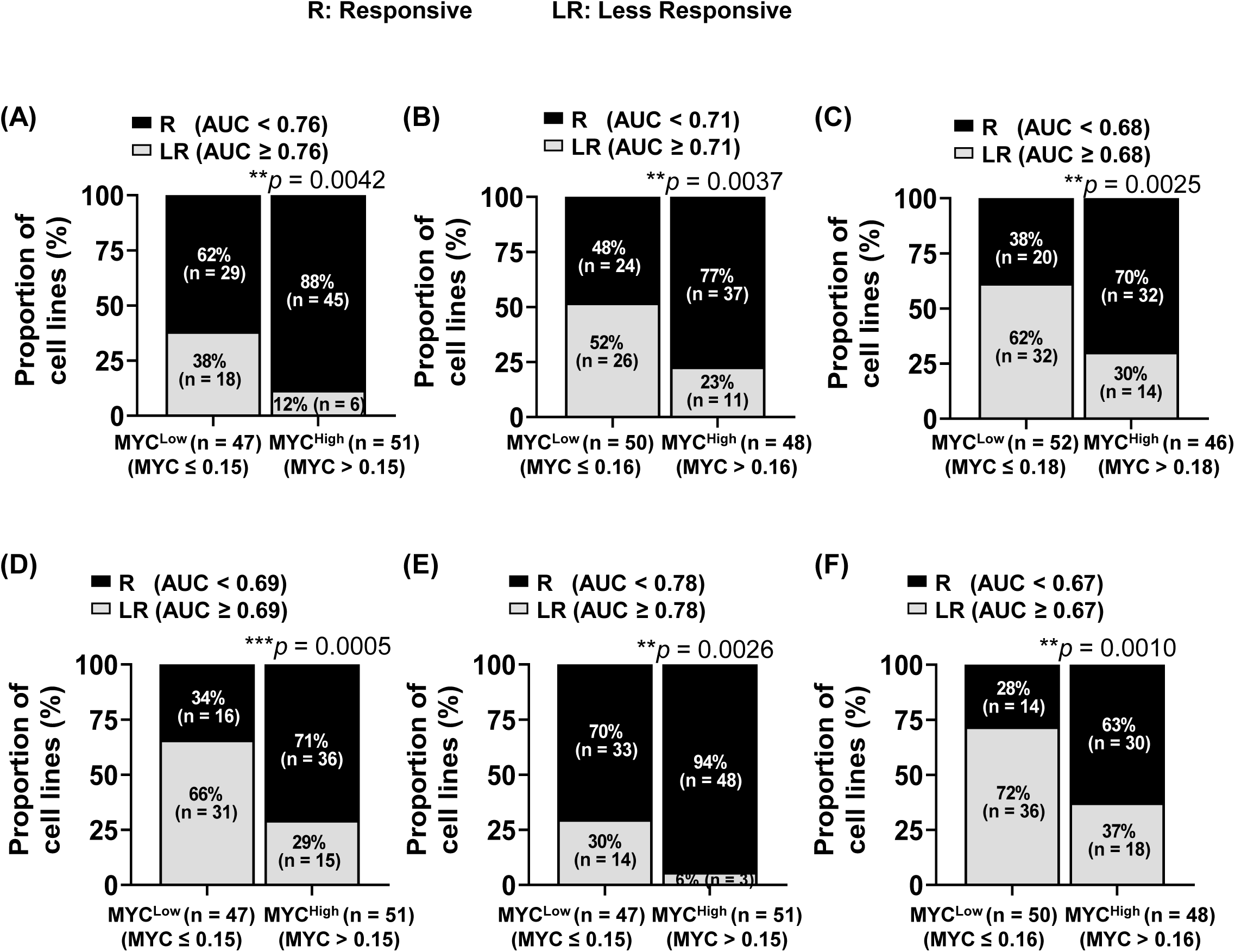
Effectiveness of MYC protein biomarker at various AUC cutoff points. R: Responsive. LR: Less Responsive. *p*-values indicate the statistical significance between the two groups. **p<0.01, ***p<0.001.

**Figure S4.**
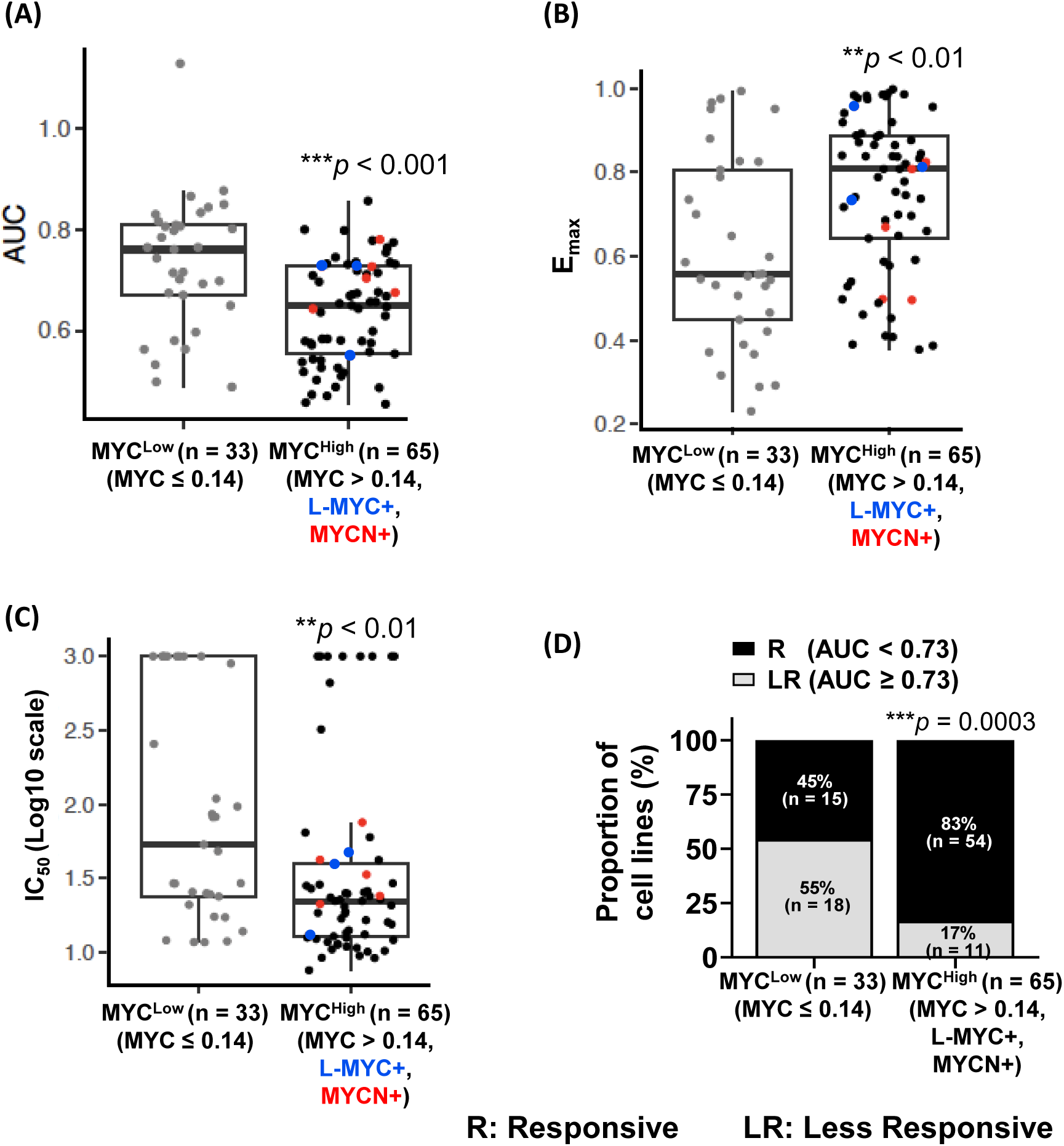
The abundance of the MYC family of oncoproteins in human cancer cell lines predicts their sensitivity to LXY18.

**Figure S5.**
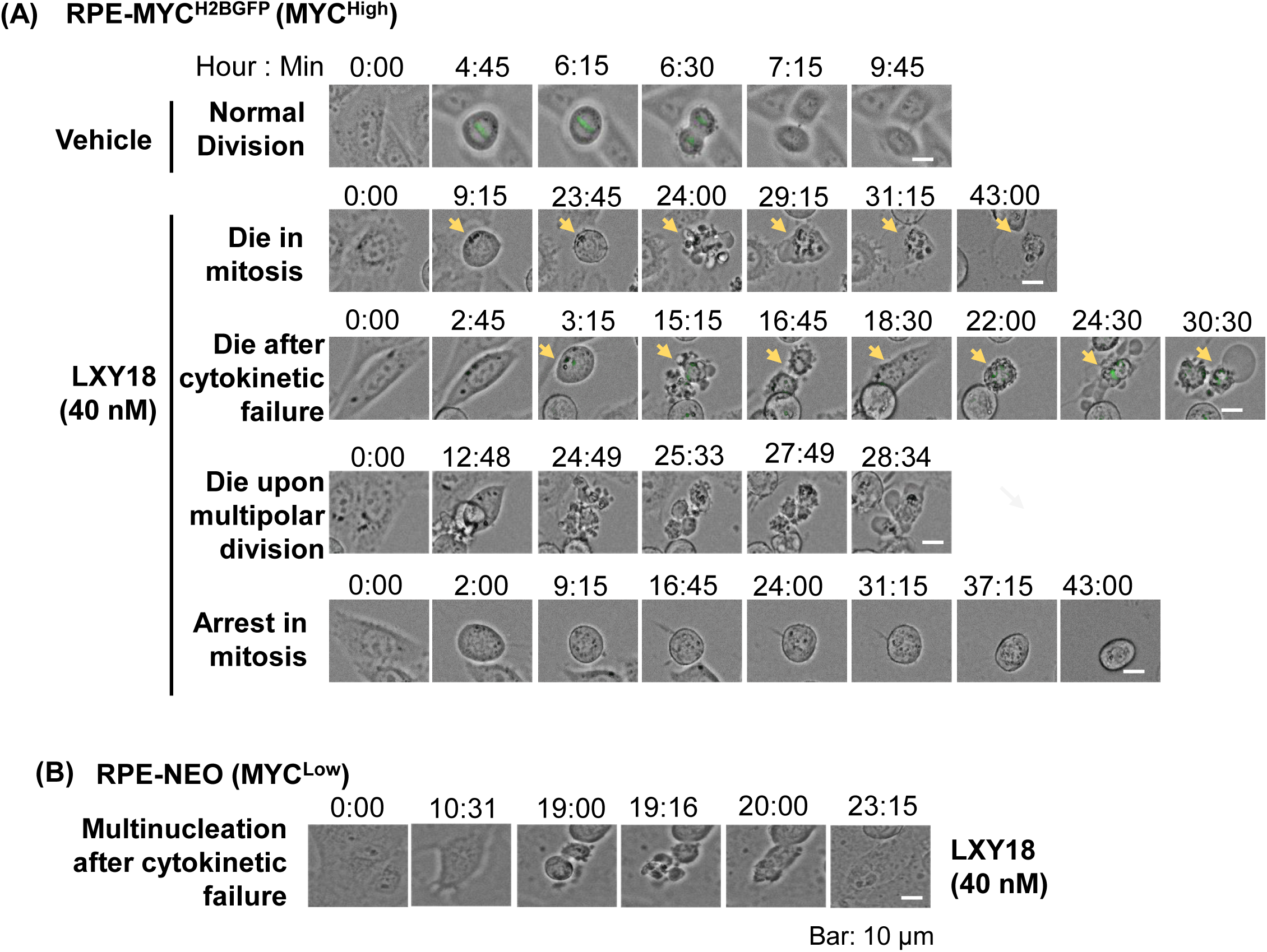
Representative time-lapse images. A. The fates of RPE-MYC^H2BGFP^ cells in the presence of 40 nM of LXY18. B. The fates of RPE-NEO cells in the presence of 40 nM of LXY18.

**Figure S6.**
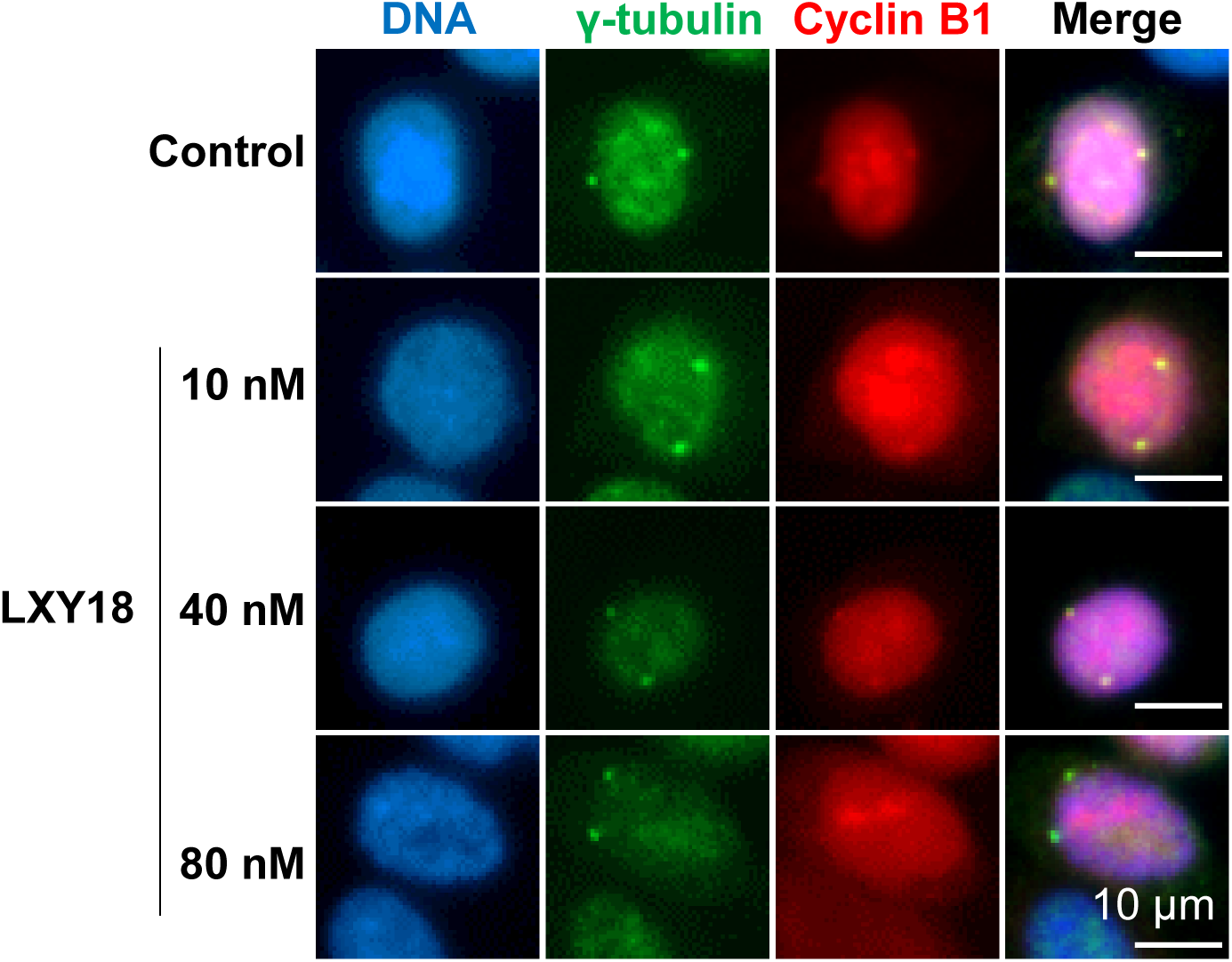
LXY18 fails to elicit excessive centrosomes in G2. NCI-H23 cells were treated with either 0.1% DMSO or indicated concentrations of LXY18 for 6 hours before fixed for double immunofluorescent staining for γ-tubulin and Cyclin B1. n > 100 in each group. Scale bar: 10 μm.

**Table S1.** A panel of 98 human cancer cell lines.

| Code | Tissue | Cell line | Code | Tissue | Cell line | Code | Tissue | Cell line |
| --- | --- | --- | --- | --- | --- | --- | --- | --- |
| 1 | Blood | Raji | 34 | Lung | HOP62 | 67 | Ovary | NCI-ADR-RES |
| 2 | Blood | Daudi | 35 | Lung | A427 | 68 | Intestine | HT-29 |
| 3 | Blood | RAMOS | 36 | Breast | BT-549 | 69 | Intestine | HCT-116 |
| 4 | Blood | U-937 | 37 | Breast | BT-474 | 70 | Intestine | SW480 |
| 5 | Blood | HL-60 | 38 | Breast | MDA-MB-157 | 71 | Intestine | HCT-15 |
| 6 | Blood | K-562 | 39 | Breast | T47D | 72 | Intestine | SW620 |
| 7 | Blood | Jurkat | 40 | Breast | MCF7 | 73 | Bone | Saos-2 |
| 8 | Blood | HPB-ALL | 41 | Breast | MDA-MB-231 | 74 | Bone | U2OS |
| 9 | Blood | MOLT-4 | 42 | Breast | MDA-MB-415 | 75 | Bone | HOS |
| 10 | Blood | CCRF-CEM | 43 | Breast | MDA-MB-436 | 76 | Cervix | Hela |
| 11 | Blood | IM-9 | 44 | Breast | MDA-MB-468 | 77 | Cervix | C-33 A |
| 12 | Blood | RPMI 8226 | 45 | Breast | MDA-MB-361 | 78 | Skin | A431 |
| 13 | Blood | NCI-BL1672 | 46 | Breast | BT-20 | 79 | Skin | UACC-257 |
| 14 | Lung | NCI-H146 | 47 | Breast | HCC1569 | 80 | Skin | SK-MEL-5 |
| 15 | Lung | NCI-H128 | 48 | Breast | ZR-75-1 | 81 | Skin | UACC-62 |
| 16 | Lung | NCI-H209 | 49 | Breast | PL-3 | 82 | Skin | SK-MEL-28 |
| 17 | Lung | NCI-H2171 | 50 | Breast | MDA-MB-461 | 83 | Skin | C32 |
| 18 | Lung | NCI-H211 | 51 | Breast | MDA-MB-175VII | 84 | Skin | MDA-MB-435S |
| 19 | Lung | NCI-H510A | 52 | Breast | SW613 | 85 | Liver | SK-Hep1 |
| 20 | Lung | NCI-H524 | 53 | Kidney | ACHN | 86 | Liver | HepG2 |
| 21 | Lung | NCI-H526 | 54 | Kidney | TUWI | 87 | Pancreas | BxPC-3 |
| 22 | Lung | NCI-H82 | 55 | Kidney | SK-N-EP1 | 88 | Pancreas | MIA Paca-2 |
| 23 | Lung | NCI-H841 | 56 | Stomach | SNU-5 | 89 | Pancreas | PANC-1 |
| 24 | Lung | NCI-H187 | 57 | Stomach | Hs746T | 90 | Pancreas | AsPC-1 |
| 25 | Lung | NCI-H446 | 58 | Stomach | NCI-N87 | 91 | Brain | AG2203 |
| 26 | Lung | NCI-H1299 | 59 | Prostate | DU-145 | 92 | Brain | SK-N-SH |
| 27 | Lung | NCI-H460 | 60 | Prostate | LNCaP | 93 | Brain | SK-N-AS |
| 28 | Lung | NCI-H23 | 61 | Prostate | VCaP | 94 | Brain | SK-N-BE2 |
| 29 | Lung | NCI-H596 | 62 | Ovary | SKOV3 | 95 | Brain | IMR-32 |
| 30 | Lung | NCI-H2170 | 63 | Ovary | OVCAR3 | 96 | Brain | T98G |
| 31 | Lung | NCI-H520 | 64 | Ovary | Caov3 | 97 | Brain | U251MG |
| 32 | Lung | A549 | 65 | Ovary | SW626 | 98 | Brain | Kelly |
| 33 | Lung | Calu6 | 66 | Ovary | PA-1 |  |  |  |

**Table S2.** Oncogenic driver mutations in human cancer cell lines.

## Material and methods

### Cell culture and chemicals

The human retinal pigment epithelial cell lines RPE-NEO, RPE-MYC, RPE- MYC^H2B-GFP^ have been described^12, 14^. The rat embryo fibroblasts Rat1A-pMig and Rat1A- MYC were also described^12^. Six human cancer cell lines, Vcap (Cat. No. SCSP-5034), Saos-2 (Cat. No. SCSP-5057), HEPG2 (Cat. No. SCSP-510), MIA Paca-2 (Cat. No. SCSP-568), PANC-1 (Cat. No. SCSP- 535), and BxPC-3 (Cat. No. TCHu 12) were purchased from National Collection of Authenticated Cell Cultures (Shanghai, China).

Four human cancer cell lines, OVCAR3 (Cat. No. CBP60294), SNU5 (Cat. No. CBP60505), Raji-Luc (Cat. No. CBP30071L), and AsPC-1-Luc (Cat. No. CBP30142L) were purchased from NanJing Cobioer Biosciences CO., LTD. (Nanjing, China). Additional 88 human cancer cell lines used in this study were purchased from American Type Culture Collection (ATCC). Under the vendor’s instruction, all cell lines were cultured in DMEM (Gibco, Cat. No. 12100061), RPMI-1640 (Gibco, Cat. No. C11875500BT), L15 (Keygen, Cat. No. KGM41300N-500), or McCoy’s 5a (Keygen, Cat. No. KGM4892N-500) supplemented with 10% fetal bovine serum (Excell, Cat. No. FSP500 10099141), penicillin (100 U/mL)-streptomycin (100 μg/mL) (Gibco, Cat. No.15140-122), 2 mM L-glutamine (Gibco, Cat. No. 25030081), and 1 mM sodium pyruvate (Gibco, Cat. No. 11360070). Cell culture was maintained in a humidified incubator with 5% CO2 at 37 °C. Mycoplasma contamination of each cell line was evaluated by using a PCR-based Myco-LumiTM Luminescent Mycoplasma Detection Kit (Beyotime, Cat. No. C0298M) and then each cell line was amplified and stored in aliquots in a liquid nitrogen tank when it was acquired. Each aliquot was used only for 10-15 passages to minimize the risk of genetic drift, ensuring consistency in the experimental data. LXY18 was synthesized in house as described^14^. Other antimitotic agents were obtained from the following commercial sources, AZD1152 (Selleck, CAS: 722544-51-6, Cat. No. S1147), AMG900 (TargetMol, CAS: 945595-80-2, Cat. No. T6380), BAY1217389 (Selleck, CAS: 1554458-53-5, Cat. No. S8215), Centrinone (TargetMol, CAS: 1798871-30-3,Cat. No. T14927), GSK923295 (TargetMol, CAS: 1088965-37-0, Cat. No. T2039), SB743921 HCl (TargetMol, CAS: 940929-33-9, Cat. No. T2255), and Sovilnesib (TargetMol, CAS: 2410796-79-9, Cat. No. T39994). RO3306 (TCL, CAS: 872573-93-8, Cat. No. R0201), Thymidine (Aladdin, CAS: 50-89-5, T104771), and MLN8237 (Selleck, CAS: 1028486-01-2, Cat. No. S1133).

### Time-lapse video recording analysis of cell division and cell death

For live-cell imaging, cells were seeded into a 12-well plate and treated with LXY18 at concentrations specified in figures 3 and S5 legend or 0.1% DMSO as vehicle control. Bright-field images were captured every 10 min for a total of 48 h beginning 5 hours after initiation of treatment using an EVOS Auto FL system equipped with an onstage incubator (ThermoFisher). The cells in mitosis were identified by their distinct morphological features, such as cell round-up, condensed chromosomes, partial alignment of chromosomes at the metaphase plate, cortex contraction, which were all readily discernible through time-lapse microscopy. The mitotic index in figure 3 was calculated by dividing the number of cells in mitosis by the total cell number in multiple randomly chosen fields. The duration of each cell spent in its first mitosis after exposed to LXY18 was also recorded. The accumulated events for the indicated fate of cells at the endpoint of this experiment were monitored and quantified. Cells in multiple randomly chosen fields in multiple wells were examined and combined data were presented in figure 3. Representative images were presented in figure S5.

### CTG and LDH assays

The cell lines were routinely passaged when reaching the mid-Log phase to avoid selection for those with less contact inhibition of proliferation. The Cell-titer Gro (CTG) assay for the live cell number and lactate dehydrogenase (LDH) assay for dead cells were utilized to evaluate effect of LXY18 on cancer and RPE model cells. Cells were routinely passaged when reaching the mid-Log phase. For the determination of the growth rate inhibition and induction of cell death, cells were plated in triplicates in 96-well plates at a density of 4,000 cells per well in 100 μL of cell culture medium. The cells were cultivated for 15∼24 h to reach a cellular confluence of ∼25 % before being exposed to LXY18 in a concentration range from 0.5 nM to 250 nM in a 2-fold of serial dilution. Cells exposed to the solvent (0.1% of DMSO) were used as a negative control. At the end of a 3-day treatment, the live cell number was quantified by the CTG assay that quantifies the ATP level (Beyotime, Cat No. C0068L). The number of dead cells was quantified by assaying LDH in cell culture medium released from dead cells using a commercial kit (Beyotime, Cat No. C0017). For the calculation of drug response descriptors, such as the half-maximal inhibitory concentration (IC_50_), the maximum effect (E_max_) and area under the response over concentration (AUC), proliferation or cell death data in the LXY18 treatment group were normalized to those in cells treated with DMSO. IC_50_ is a descriptor for drug potency without reflecting an effect size, whereas Emax is a parameter for efficacy. AUC combines potency and efficacy into a single parameter, which can be conveniently and more accurately compared for a single drug across multiple cell lines exposed to the same range of drug concentrations. The normalized data were then fit to the four parameters logistic curve using GraphPad Prism 8.0.2. Statistical analysis was performed using the same software. Data are presented as mean ± SD.

### Trypan blue exclusion assay

The Trypan blue exclusion assay uses Trypan blue dye to selectively stain dead cells. Cells were harvested by trypsinization before suspended in sterile PBS. A 0.4% of Trypan blue dye solution (Cat. NO. E607338, Sangon, Shanghai, China) was added to the cell suspension in a 1:1 ratio (v/v) and the cell suspension was immediately loaded into a Hemocytometer chamber (XB-K-25, SMIC). Under an inverted tissue culture microscopy, cells positive or negative for Trypan blue-staining were enumerated to indicate dead and live cells, respectively. The fraction of dead cells was calculated by dividing the number of dead cells by the total number of cells.

### Cell synchronization

Cells were synchronized at the G1/S boundary by exposure to 2 .5 mM of thymidine for 24 h before released for 16 h into a thymidine-free medium containing 5 μM of CDK1 inhibitor RO3306 that arrested cells at the G2/M transition. The cells were then released into a drug-free medium containing different mitotic inhibitors for 30 min in Figure 6. Or cells were released into a fresh drug-free medium for 50 min, then exposed to the drugs 10 min or 30min in Figure 7 before being fixed for immunofluorescent analysis.

### Western blot analysis

The RIPA buffer (50mM Tris, 150mM NaCl, 1% TritonX-100, 1% sodium deoxycholate, 0.1%SDS) supplemented with a cocktail of phosphatase (Beyotime, #P1082) and protease inhibitors (Beyotime, # P1005) was used to lyse cells on ice for 15 min in 1.5 ml Eppendorf tubes. The cells were then sonicated for 10 seconds before centrifuged at 12,000 rpm for 15 min. The protein concentration in lysate was quantified by using the BCA assay kit (Sangon Biotech, Cat. No. C503051-0500) according to the manufacturer’s instructions. After boiling in an SDS-PAGE sample loading buffer, ∼50 μg of proteins were loaded into each lane of a precasted protein gel (Invitrogen, NP0336BOX or NP0321BOX), were separated under a voltage of 100-130V for 1∼2h and were transferred onto a PVDF membrane (0.22 μm) for 80 min at a constant voltage of 70V. The membrane was blocked for 1.5 h with 5% nonfat milk (Sangon, A600669-0250) in a Tris-buffered saline with 0.1% Tween (TBST) at room temperature before incubation overnight at 4 °C with a primary antibody in an Odyssey blocking buffer or in phosphate buffered saline (PBS) containing 0.2% Tween 20). The primary and secondary antibodies used are at below. A rabbit anti-c-Myc mAb (Y69) was from Abcam (ab32072) and used a 1:1000 dilution. A rabbit anti-N-myc (ab24193) was from Abcam and used a 1:1000 dilution. A rabbit anti-L-myc (76266S) was from CST and used a 1:1000 dilution. A mouse anti-actin antibody (Cat. #. 66009-1-1g) was from Proteintech and was used at a 1:10000 dilution. The membrane was incubated with a secondary antibody at room temperature for 1 h. An IRDye® 800CW Goat anti-Mouse IgG (Cat. #. 926-32210, Lot#C91210-09) and an IRDye® 680RD Goat anti-Rabbit antibody (926-68071, Lot#D00115-06) were used at a 1:10000 and 1:5000 dilution, respectively. Images were captured using an Odyssey CLx Infrared Imaging System with over 6 logs of linear dynamic range for accurate quantification and processed with Image Studio Ver 5.2. For easy comparison of MYC abundance among different cell lines, the relative abundance of MYC was calculated by normalizing MYC in a cancer cell line to that in LXY18-sensitive model cell line RPE-MYC on the same membrane. The abundance of MYC in RPE-MYC cell line was set as 1.0.

### Immunofluorescent analysis

Cells were cultured on glass cover slips, fixed in 4% paraformaldehyde (PFA), and permeabilized with 0.1% Triton X-100 before incubating with antibodies against H3Ser10P (CST, Cat. No. 53348). β-tubulin (Sigma, Cat. No. T5201), and AURKA Thr288P (CST, Cat. No. 3079), γ-tubulin (Proteintech, 66320-1-Ig), and CyclinB 1(Santa Cruz, SC-752). After staining with an appropriate secondary antibody, cells were mounted with DAPI-containing mounting solution and imaged under an EVOS FL auto microscope. Secondary antibodies Fluorescein (FITC)-conjugated Affinipure Goat Anti-Rabbit IgG(H+L) (Cat. No. 111-095-003) and Rhodamine (TRITC) AffiniPure Goat Anti-Mouse IgG (H+L) (Cat. No. 115-025-003) were purchased from Jackson.

## Conceptual Abstract

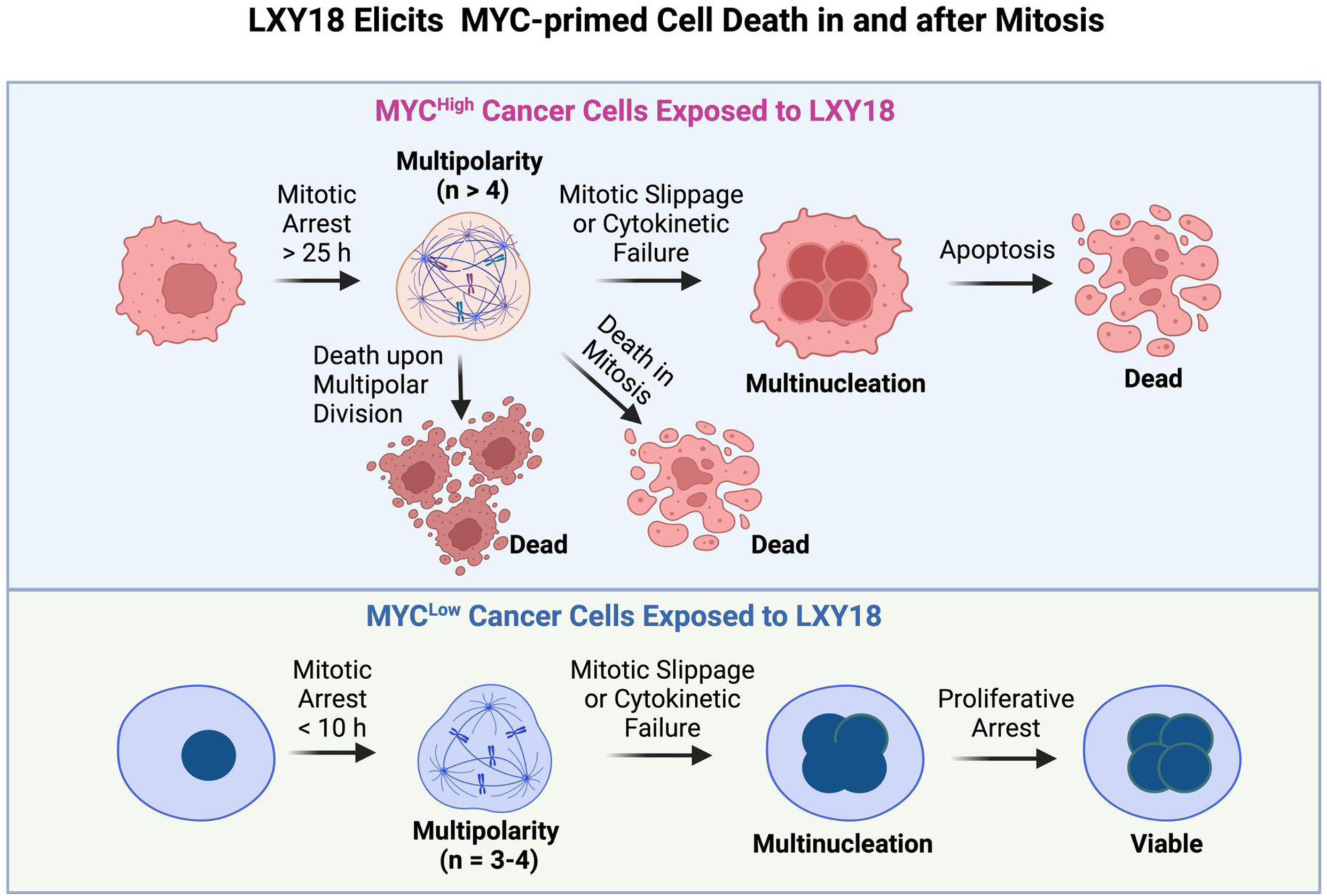

