## Supplementary information file for "MYC amplifies mitotic perturbations elicited by LXY18 to enable synthetic lethality"

91 cell lines

| DepMap_ID | Name |
|---|---|
| ACH-000757 | A427 |
| ACH-001328 | A431 |
| ACH-000681 | A549 |
| ACH-000046 | ACHN |
| ACH-000222 | ASPC1 |
| ACH-000536 | BT20 |
| ACH-000927 | BT474 |
| ACH-000288 | BT549 |
| ACH-000535 | BXPC3 |
| ACH-000580 | C32 |
| ACH-001333 | C33A |
| ACH-000264 | CALU6 |
| ACH-000713 | CAOV3 |
| ACH-001738 | CCRFCEM |
| ACH-000786 | DAUDI |
| ACH-000979 | DU145 |
| ACH-000930 | HCC1569 |
| ACH-000971 | HCT116 |
| ACH-000997 | HCT15 |
| ACH-001086 | HELA |
| ACH-000739 | HEPG2 |
| ACH-000002 | HL60 |
| ACH-000861 | HOP62 |
| ACH-000613 | HOS |
| ACH-000942 | HPBALL |
| ACH-000616 | HS746T |
| ACH-000552 | HT29 |
| ACH-002247 | IM9 |
| ACH-000310 | IMR32 |
| ACH-000995 | JURKAT |
| ACH-000551 | K562 |
| ACH-000259 | KELLY |
| ACH-000977 | LNCAP |
| ACH-000019 | MCF7 |
| ACH-000621 | MDAMB157 |
| ACH-000759 | MDAMB175VII |
| ACH-000768 | MDAMB231 |
| ACH-000934 | MDAMB361 |
| ACH-000876 | MDAMB415 |
| ACH-000884 | MDAMB435S |
| ACH-000573 | MDAMB436 |
| ACH-000849 | MDAMB468 |
| ACH-000601 | MIAPACA2 |
| ACH-001127 | MOLT4 |
| ACH-002286 | NCIH128 |
| ACH-000510 | NCIH1299 |
| ACH-000506 | NCIH146 |
| ACH-001136 | NCIH187 |

| | |
|---|---|
| ACH-000290 | NCIH209 |
| ACH-000639 | NCIH211 |
| ACH-000481 | NCIH2170 |
| ACH-000525 | NCIH2171 |
| ACH-000900 | NCIH23 |
| ACH-000800 | NCIH446 |
| ACH-000463 | NCIH460 |
| ACH-000871 | NCIH510A |
| ACH-000395 | NCIH520 |
| ACH-000816 | NCIH524 |
| ACH-000767 | NCIH526 |
| ACH-000628 | NCIH596 |
| ACH-000355 | NCIH82 |
| ACH-000292 | NCIH841 |
| ACH-000427 | NCIN87 |
| ACH-000001 | NIHOVCAR3 |
| ACH-001374 | PA1 |
| ACH-000164 | PANC1 |
| ACH-000654 | RAJI |
| ACH-001636 | RAMOS |
| ACH-000817 | RPMI8226 |
| ACH-000410 | SAOS2 |
| ACH-000361 | SKHEP1 |
| ACH-000615 | SKMEL28 |
| ACH-000730 | SKMEL5 |
| ACH-000260 | SKNAS |
| ACH-000312 | SKNBE2 |
| ACH-001192 | SKNEP1 |
| ACH-000149 | SKNSH |
| ACH-000811 | SKOV3 |
| ACH-000303 | SNU5 |
| ACH-000842 | SW480 |
| ACH-000651 | SW620 |
| ACH-001399 | SW626 |
| ACH-000147 | T47D |
| ACH-000571 | T98G |
| ACH-000232 | U251MG |
| ACH-000364 | U2OS |
| ACH-000406 | U937 |
| ACH-000579 | UACC257 |
| ACH-000425 | UACC62 |
| ACH-000115 | VCAP |
| ACH-000097 | ZR751 |

*RAS*-*RAF* mutant

| DepMap_ID | Name |
|---|---|
| ACH-000002 | HL60 |
| ACH-000097 | ZR751 |
| ACH-000149 | SKNSH |
| ACH-000164 | PANC1 |
| ACH-000222 | ASPC1 |
| ACH-000260 | SKNAS |
| ACH-000361 | SKHEP1 |
| ACH-000425 | UACC62 |
| ACH-000463 | NCIH460 |
| ACH-000510 | NCIH1299 |
| ACH-000552 | HT29 |
| ACH-000579 | UACC257 |
| ACH-000580 | C32 |
| ACH-000601 | MIAPACA2 |
| ACH-000615 | SKMEL28 |
| ACH-000651 | SW620 |
| ACH-000681 | A549 |
| ACH-000730 | SKMEL5 |
| ACH-000739 | HEPG2 |
| ACH-000757 | A427 |
| ACH-000768 | MDAMB231 |
| ACH-000817 | RPMI8226 |
| ACH-000842 | SW480 |
| ACH-000861 | HOP62 |
| ACH-000884 | MDAMB435S |
| ACH-000900 | NCIH23 |
| ACH-000934 | MDAMB361 |
| ACH-000971 | HCT116 |
| ACH-000997 | HCT15 |
| ACH-001127 | MOLT4 |
| ACH-001374 | PA1 |
| ACH-001399 | SW626 |
| ACH-001738 | CCRFCEM |
| ACH-002247 | IM9 |

*RB1* mutant

| DepMap_ID | Name |
| --- | --- |
| ACH-000290 | NCIH209 |
| ACH-000506 | NCIH146 |
| ACH-000525 | NCIH2171 |
| ACH-000536 | BT20 |
| ACH-000786 | DAUDI |
| ACH-000816 | NCIH524 |
| ACH-000979 | DU145 |
| ACH-000995 | JURKAT |

*PTEN -PI3K* mutant

| DepMap_ID | Name |
|---|---|
| ACH-000019 | MCF7 |
| ACH-000097 | ZR751 |
| ACH-000147 | T47D |
| ACH-000463 | NCIH460 |
| ACH-000536 | BT20 |
| ACH-000552 | HT29 |
| ACH-000571 | T98G |
| ACH-000580 | C32 |
| ACH-000615 | SKMEL28 |
| ACH-000628 | NCIH596 |
| ACH-000730 | SKMEL5 |
| ACH-000811 | SKOV3 |
| ACH-000876 | MDAMB415 |
| ACH-000927 | BT474 |
| ACH-000934 | MDAMB361 |
| ACH-000971 | HCT116 |
| ACH-000995 | JURKAT |
| ACH-000997 | HCT15 |
| ACH-001333 | C33A |
| ACH-001738 | CCRFCEM |

*TP53* mutant

| DepMap_ID | Name |
|---|---|
| ACH-000115 | VCAP |
| ACH-000147 | T47D |
| ACH-000164 | PANC1 |
| ACH-000232 | U251MG |
| ACH-000259 | KELLY |
| ACH-000264 | CALU6 |
| ACH-000288 | BT549 |
| ACH-000292 | NCIH841 |
| ACH-000312 | SKNBE2 |
| ACH-000355 | NCIH82 |
| ACH-000395 | NCIH520 |
| ACH-000481 | NCIH2170 |
| ACH-000525 | NCIH2171 |
| ACH-000535 | BXPC3 |
| ACH-000536 | BT20 |
| ACH-000552 | HT29 |
| ACH-000571 | T98G |
| ACH-000601 | MIAPACA2 |
| ACH-000613 | HOS |
| ACH-000615 | SKMEL28 |
| ACH-000616 | HS746T |
| ACH-000628 | NCIH596 |
| ACH-000651 | SW620 |
| ACH-000654 | RAJI |
| ACH-000713 | CAOV3 |
| ACH-000768 | MDAMB231 |
| ACH-000786 | DAUDI |
| ACH-000800 | NCIH446 |
| ACH-000816 | NCIH524 |
| ACH-000817 | RPMI8226 |
| ACH-000842 | SW480 |
| ACH-000849 | MDAMB468 |
| ACH-000871 | NCIH510A |
| ACH-000876 | MDAMB415 |
| ACH-000884 | MDAMB435S |
| ACH-000900 | NCIH23 |
| ACH-000927 | BT474 |
| ACH-000930 | HCC1569 |
| ACH-000934 | MDAMB361 |
| ACH-000942 | HPBALL |
| ACH-000979 | DU145 |
| ACH-000995 | JURKAT |
| ACH-000997 | HCT15 |
| ACH-001127 | MOLT4 |
| ACH-001136 | NCIH187 |
| ACH-001328 | A431 |
| ACH-001333 | C33A |
| ACH-001399 | SW626 |

ACH-001636  RAMOS
ACH-001738  CCRFCEM
ACH-002286  NCIH128

*MYC* family amplification

| Cell Line Name | *L-MYC* | *MYCN* | *MYC* |
|---|---|---|---|
| A549 | | | |
| ACHN | | | |
| ASPC1 | | | |
| BT20 | | | |
| BT474 | | | |
| BT549 | | | Amplified |
| BXPC3 | | | |
| C32 | | | |
| CALU6 | | | |
| CAOV3 | | | |
| DAUDI | | | |
| DU145 | | | |
| HCC1569 | | | Amplified |
| HCT116 | | | |
| HCT15 | | | |
| HEPG2 | | | |
| HL60 | | | Amplified |
| HOP62 | | | |
| HOS | | | |
| HPBALL | | | |
| HS746T | | | |
| HT29 | | | Amplified |
| IMR32 | | Amplified | |
| JURKAT | | | |
| K562 | | | |
| KELLY | | Amplified | |
| LNCAP | | | |
| MCF7 | | | Amplified |
| MDAMB157 | | | Amplified |
| MDAMB175VII | | | |
| MDAMB231 | | | |
| MDAMB361 | | | |
| MDAMB415 | | | |
| MDAMB435S | | | |
| MDAMB436 | | | Amplified |
| MDAMB468 | Amplified | | |
| MIAPACA2 | | | |
| MOLT4 | | | |
| NCIH1299 | | | |
| NCIH146 | | | |
| NCIH209 | Amplified | | |
| NCIH211 | | | Amplified |
| NCIH2170 | | | Amplified |
| NCIH2171 | | | Amplified |
| NCIH23 | | | Amplified |
| NCIH446 | | | Amplified |
| NCIH460 | | | Amplified |
| NCIH510A | Amplified | | Amplified |

| | | | |
|---|---|---|---|
| NCIH520 | Amplified | | |
| NCIH524 | | | Amplified |
| NCIH526 | Amplified | Amplified | |
| NCIH596 | | | |
| NCIH82 | | | Amplified |
| NCIH841 | | | |
| NCIN87 | | | Amplified |
| NIHOVCAR3 | | | |
| PANC1 | | | |
| RAJI | | | |
| RPMI8226 | | | |
| SKHEP1 | | | |
| SKMEL28 | | | |
| SKMEL5 | | | |
| SKNAS | | | |
| SKNBE2 | | Amplified | |
| SKNSH | | | |
| SKOV3 | | | |
| SNU5 | | | |
| SW480 | | | Amplified |
| SW620 | | | Amplified |
| T47D | | | |
| T98G | | | |
| U251MG | | | |
| U2OS | | | Amplified |
| U937 | | | |
| UACC257 | | | |
| UACC62 | | | |
| VCAP | | | Amplified |
| ZR751 | | | |
